# Distinct spatial maps and multiple object codes in the lateral entorhinal cortex

**DOI:** 10.1101/2022.12.19.521018

**Authors:** Xu Huang, Magdalene Isabell Schlesiger, Isabel Barriuso-Ortega, Christian Leibold, Duncan Archibald Allan MacLaren, Nina Bieber, Hannah Monyer

**Affiliations:** Department of Clinical Neurobiology at the Medical Faculty of Heidelberg University and German Cancer Research Center (DKFZ), Im Neuenheimer Feld 280, 69120 Heidelberg, Germany; Institute Biology III & Bernstein Center Freiburg, University of Freiburg, Germany; BrainLinks-BrainTools, University of Freiburg, Germany

## Abstract

The lateral entorhinal cortex (LEC) is a major cortical input area to the hippocampus and it is crucial for associative object-place-context memories. An unresolved question is whether these associations are performed exclusively in the hippocampus or also upstream thereof. Anatomical evidence suggests that the LEC processes both object and spatial information. We here describe a gradient of spatial selectivity along the antero-posterior axis of the LEC. We demonstrate that the LEC generates distinct spatial maps for different contexts that are independent of object coding and vice versa, thus providing evidence for pure spatial and pure object codes upstream of the hippocampus. Whilst space and object coding occur by and large separately in the LEC, we identified neurons that encode for space and objects conjunctively. Together, these findings point to a scenario in which the LEC sustains both distinct space and object coding as well as associative space-object coding.

## Introduction

The hippocampus and its interaction with many neocortical areas subserve the integration of spatial and non-spatial information in a temporally-ordered manner, which is crucial for the formation of episodic memories^1–5^. The entorhinal cortex, comprising the medial and lateral subdivisions (MEC and LEC), is the main interface between neocortical areas and the hippocampus^6,7^. Spatial and non-spatial information are processed in visual associative areas by the dorsal and ventral streams, respectively, which convey this information to the entorhinal cortex via the postrhinal (dorsal stream) and perirhinal cortex (ventral stream)^8–12^. The MEC receives major cortical input from the postrhinal cortex and is specialized in spatial coding^13,14^, as evidenced by the existence of a plethora of spatially modulated neurons^15–18^. The LEC, on the other hand, receives input from the perirhinal cortex and was long considered a non-spatial region^13,14^. Specifically, LEC neurons were reported to encode non-spatial features, such as objects, odors and time^19–22^. In addition, LEC neurons were shown to signal the animal’s position related to the environmental boundaries and/or objects^23^. Notably, in contrast to MEC neurons, LEC neurons did not exhibit spatial selectivity in empty open field environments^24,25^. However, a recent anatomical study demonstrated that the LEC receives input both from perirhinal and postrhinal cortices to a similar extent^26^. This raises the question to which extent LEC neurons support spatial coding. Indeed, it was found that, in addition to object coding, LEC neurons displayed increased spatial selectivity in the presence of objects^19,20,27^. In yet another study, a small proportion of LEC neurons developed spatial firing fields in places where objects had previously been^28^.

At the behavioral level, the LEC was shown to be required for the formation of memories that involve the association of object and spatial context information^29–31^. The joint targeting of LEC neurons by perirhinal and postrhinal projections^26^ raises furthermore the question of whether and, if so, how spatial and object information is integrated at the cellular level. Here, we set out to bridge the gap between anatomical and behavioral data by addressing this question at the single-neuron level in mice exploring different combinations of spatial contexts and objects.

We systematically investigated spatial coding in the LEC and searched for the presence of a putative gradient, as was shown for other areas. For instance, in the hippocampus and the MEC, there is a dorsoventral gradient that determines the field size of spatially modulated neurons^32–35^. Notably, we found a spatial selectivity gradient along the antero-posterior axis in the LEC. Spatially modulated neurons were apparent when the animal foraged in an empty open field and remapped across spatial contexts. These findings prompted us to further investigate whether space and object coding is performed by discrete populations of neurons or whether individual neurons represent space and objects conjunctively. In other words, we addressed the question whether, upstream of the hippocampus, there is an object representation that is independent from the spatial representation, as predicted by the cognitive map theory^36–38^, or whether joint information processing occurs already at this level, as suggested by other anatomy-based theories^39,40^.

## Results

### Spatial selectivity of neurons increases from posterior to anterior LEC

We first analyzed the spatial properties of neurons in the anterior (A), intermediate (I) and posterior (P) LEC. For unambiguous identification of the tetrode locations, we implanted 5 mice with a three-tetrode microdrive arranged such that each tetrode would target distinct anteroposterior positions in the LEC (Figure 1A, see Figure 1B and Figure S1 for histology with tetrode locations). LEC neuronal activity was recorded while mice foraged for food in at least two 20-min open field trials. There was a gradual increase of the spatial information scores of putative excitatory neurons (mean firing rate > 0.1 and < 5 Hz, spike waveform asymmetry < 0.1 and spike trough-to-peak latency > 0.4 ms) from posterior to more intermediate and anterior locations (median: A: 0.21; I: 0.16; P: 0.10) (Figure 1C). Of note, there was a spatial selectivity gradient in all animals (Figure S1), and the gradual decrease in spatial information was paralleled by an increase in the number of firing fields (median: A: 2.00; I: 4.00; P: 6.00) (Figures 1D and 1E). The finding of discernible, yet not well delineated, spatial firing fields prompted the question of how anterior/intermediate LEC spatial firing compares to that of MEC and CA1.

**Figure 1.**
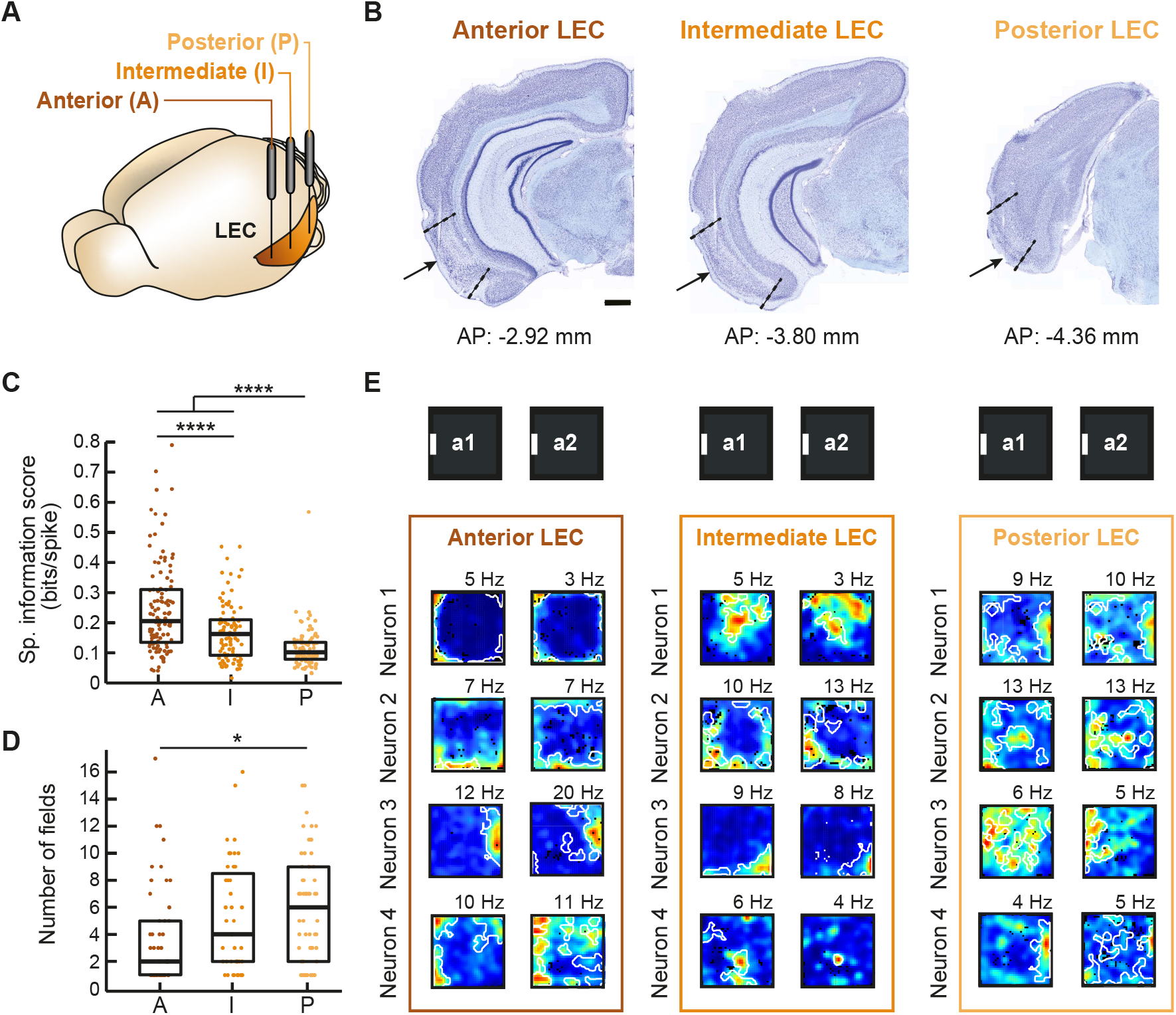
Gradient of spatial selectivity in LEC neurons along the antero-posterior axis. **(A)** Scheme depicting the mouse brain and the LEC. In each animal, a microdrive with three tetrodes was used to target LEC superficial layers at anterior (A), intermediate (I) and posterior (P) positions. **(B)** Cresyl violet-stained coronal sections from a representative mouse. The position of the three tetrode tips is indicated by an arrow. The antero-posterior (AP) coordinates from bregma are indicated below. See Figure S1 for histology from the other mice. **(C)** Spatial information scores of all excitatory neurons recorded in each of the LEC anteroposterior locations (A: anterior, I: intermediate, P: posterior). Scores are significantly different across the different LEC antero-posterior positions (p < 0.0001, H = 63.726, df = 2, Kruskal-Wallis ANOVA, anterior LEC: n = 113 neurons from 4 mice; intermediate LEC: n = 92 neurons from 5 mice; posterior LEC: n = 90 neurons from 5 mice). **(D)** Number of firing fields increases from anterior to posterior LEC positions (p < 0.05, H = 8.7103, df = 2, Kruskal-Wallis ANOVA). **(F)** Color-coded firing rate maps of representative neurons recorded at each of the three LEC antero-posterior positions while mice foraged in a familiar black, square open field environment with a white cue card on one of the walls. Firing fields are delineated in white in each map. Four neurons recorded in two consecutive 20-min sessions (a1 and a2) are shown per location. Maps are scaled from dark blue (locations where the neuron was silent) to red (peak rate, indicated on top of each map in Hz). Pixels not sampled are black. Boxplots show medians and interquartile ranges, and each point represents an individual neuron. Holm-Bonferroni correction procedure was applied for multiple comparisons. *p<0.05, ****p < 0.0001.

### Superficial layers of the anterior/intermediate LEC harbor a high proportion of spatially modulated neurons

We recorded the activity from neurons predominantly in the superficial layers of the anterior/intermediate LEC of 5 mice (Figure S2). Many neurons displayed firing fields that were stable across trials (Figure 2A). We next compared their firing properties to those of neurons in the hippocampal CA1 region, MEC/parasubiculum (referred to as MEC) (Figures 2B and 2C), and to non-fast-spiking neurons in the medial septum (MS) (Figure 2D) which have low, if any, spatial selectivity^41^. LEC neurons exhibited significantly higher spatial information scores compared to MS neurons, but lower spatial information scores than those of MEC and CA1 neurons (median: LEC, 0.24; MEC, 0.58; CA1, 0.96; MS, 0.06) (Figure 2E). Similar region-dependent differences were apparent when the within-trial map stability comparing the firing rate maps of the 1^st^ with the 2^nd^ half of the open field trial was assessed (median: LEC, 0.36; MEC, 0.71; CA1, 0.67; MS, 0.14) (Figure 2E). Finally, when restricting the analysis to the last recording session per mouse (ascertaining that the recordings stemmed from histologically confirmed locations of the tetrode tips in layer 2), the values were comparable to those obtained when considering the entire LEC dataset (median spatial information score: 0.22; within-trial map stability: 0.40; n = 38 neurons in 5 mice). To determine the proportion of spatially modulated neurons in each brain region, we compared the spatial information score and within-trial map stability with the one that would be expected by chance for each individual neuron (Figure 2F, see ref.^42^ for a comparable classification method). The analysis revealed that the majority of putative excitatory LEC neurons were spatial (67.8%), and again, the proportion of spatially modulated neurons in the LEC was lower than in the MEC and CA1 (97.0% and 91.8%, respectively), but considerably higher than in the MS (20.5%) (Figure 2G).

**Figure 2.**
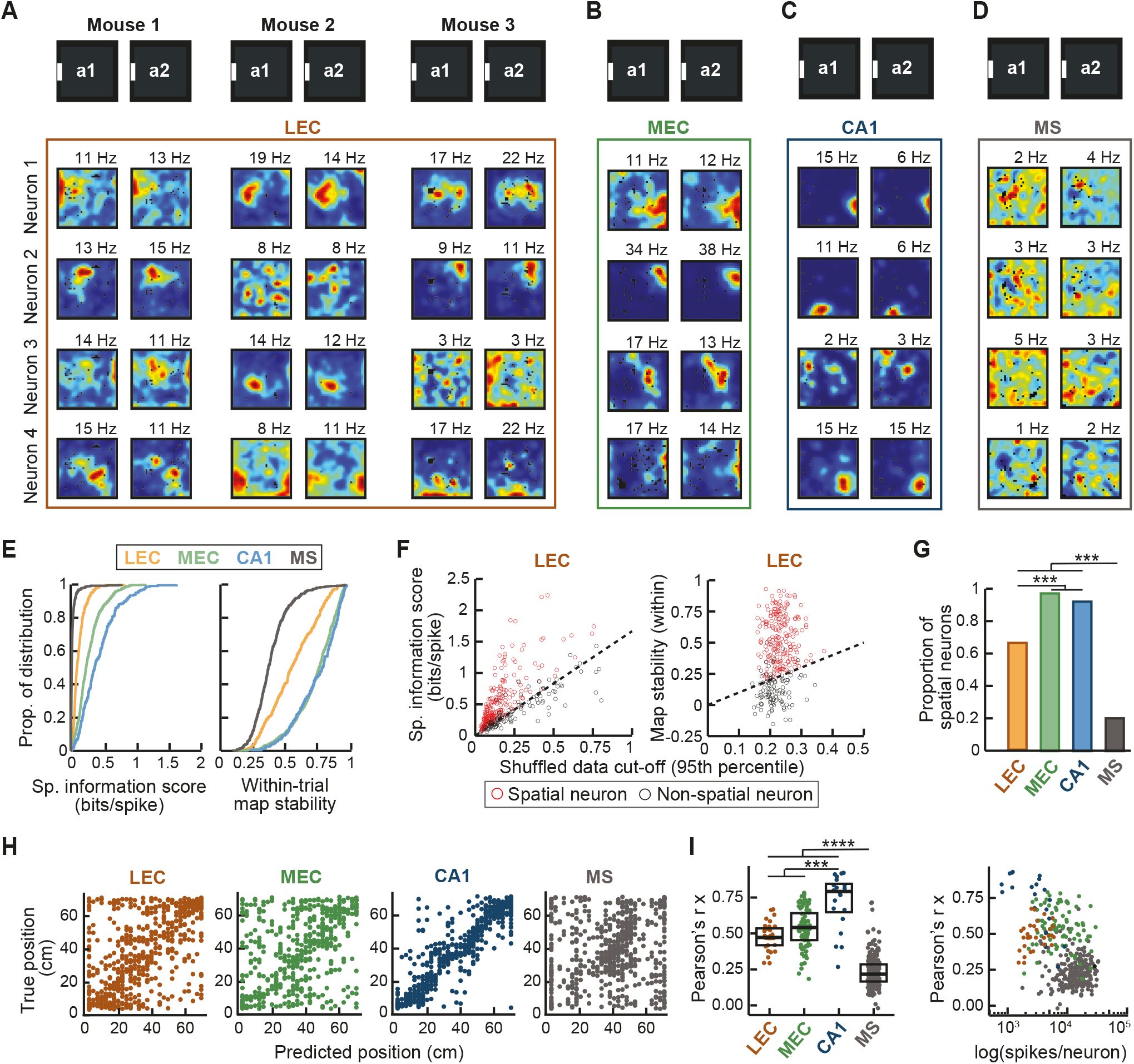
Spatial features of LEC, MEC, CA1 and MS neurons. **(A)** Spatial firing characteristics of neurons in the anterior/intermediate LEC. See Figure S2 for histological examination of recording locations. Colour-coded firing rate maps of representative LEC neurons recorded in three mice foraging in a familiar black, square open field environment with a white cue card. Four neurons recorded simultaneously in two consecutive 20-min sessions (a1 and a2) are shown per mouse. Maps are scaled from dark blue (locations where the neuron was silent) to red (peak rate, indicated on top of each map in Hz). Pixels not sampled are black. **(B-D)** Rate maps of neurons recorded in MEC **(B)**, hippocampal area CA1 **(C)** and the MS **(D). (E)** Spatial information scores (left) and within-trial map stability (right) in the entire population of putative excitatory neurons in the four brain areas. The spatial information score and within-trial map stability is highest in hippocampal neurons (blue), followed by MEC (green), LEC (brown) and MS (grey) neurons (all p values < 0.001, H = 748.84 and H = 540.35, respectively, df = 3, Kruskal-Wallis ANOVA, n neurons in m mice, LEC, 211 and 5, MEC, 298 and 10, CA1, 240 and 4, MS, 375 and 4). **(F)** For each LEC neuron, the spatial information score (left) and within-trial map stability (right) is plotted against the corresponding value that would be expected by chance (95^th^ percentile cut-off, see STAR Methods). **(G)** In the LEC, the proportion of spatially modulated neurons was higher than in the MS but lower than in the hippocampus and MEC (all p values < 0.001, X-squared = 531.66, df = 3, χ^2^ test). **(H)** Decoding of the position of the animal via minimum mean square estimator. Place decoding based on the activity of LEC, MEC, CA1 and MS neurons in one example session for each region. Scatter plots display the estimated x position versus the true x or y position of the animal as decoded by the activity of simultaneously recorded neurons in each region. Only sessions with more than 10 co-recorded neurons were considered for this analysis. **(I)** Median Pearson correlation (r) of the fitted x coordinate calculated per session for the four areas. True x position is highly correlated to the estimated position based on CA1 population activity. The estimations derived from MEC and LEC activity are similar, and they are both lower than that from CA1. MS population activity does not encode for position (all p-values < 0.001, H = 218.06, df = 3, Kruskal-Wallis ANOVA, LEC: 32 sessions; MEC: 88 sessions; CA1: 20 sessions; MS: 228 sessions). **(J)** Median Pearson correlation (r) of the fitted x coordinate as a function of the mean spikes per neuron for each session. MEC neurons exhibit a higher number of spikes per session as compared to those in LEC, but the Pearson correlation is not significantly different (p < 0.0001, Kruskal-Wallis ANOVA). Boxplots show medians and interquartile ranges, and each point represents one session. Holm-Bonferroni correction procedure was applied for multiple comparisons. ***p < 0.001, ****p <0.0001.

We also compared how well the position of the animal could be predicted based on the activity of simultaneously recorded cells in either the LEC, MEC, CA1 or MS. The reconstructed x and y coordinates within the open field estimated from LEC neuron activity correlated with the true position of the mouse (median Pearson correlation coefficient (r) for estimated vs. true x and y coordinates: 0.47 and 0.36) (Figures 2H and 2I). The cue card was located in the y axis, ruling out the possibility that the better estimation along the x axis is driven by the representation of the cue card. The estimation along the x axis was comparable for LEC and MEC neuron activity (r for x: 0.54; r for y: 0.58), but it was better along the y axis for MEC than for LEC (Kruskal-Wallis test, H = 200.63, all post-hoc comparisons p<0.0001). The best estimation could be derived from CA1 neuronal activity (0.79 and 0.80), and the worst was obtained from the MS population (0.21 and 0.22). This result held up irrespective of the number of simultaneously recorded neurons (Figure S2C,D). Notably, the average number of spikes per neuron and session was lower for LEC than for MEC (median: LEC, 2595 spikes; MEC, 7508 spikes), although the correlation between estimated and true position was similar (Figure 2J and Figure S2E).

### A fraction of fast-spiking neurons are spatially modulated

It has been established that the precise tuning of principal neurons is governed by fast-spiking (FS) neurons in the local network in many cortical regions^43–46^. Furthermore, there is evidence that FS neurons in the MEC carry spatial information to some extent^47^. We hence compared the spatial firing of FS neurons (mean firing rate > 5 Hz, spike waveform asymmetry > 0.1, spike trough-to-peak latency < 0.4 ms) in the four regions. In agreement with anatomical studies where FS neurons were identified by parvalbumin immunoreactivity^48^, there was a lower proportion of FS neurons in LEC (2.4%) than in MEC (11.6%) (Figures 3A and 3B). The proportion of FS neurons in CA1 and MS was 6.1% and 23.0%, respectively. The mean firing rate was not different for FS neurons in the four brain areas (median: LEC, 20.8 Hz; MEC, 21.7 Hz; CA1, 22.8 Hz; MS, 18.9 Hz) (Figures 3C and 3D). Notably, 23.1-55.6% of FS neurons in the three hippocampal-parahippocampal regions, but only 6.5% of FS neurons in the MS, were spatially modulated (Figure 3E). Importantly, the firing of spatial FS neurons remained stable between consecutive open field trials (Figure 3F, median across-trial map stability: LEC, 0.70; MEC, 0.60; CA1, 0.79; MS, 0.49), suggesting that FS neurons are part of the mapping system in the hippocampal-parahippocampal regions.

**Figure 3.**
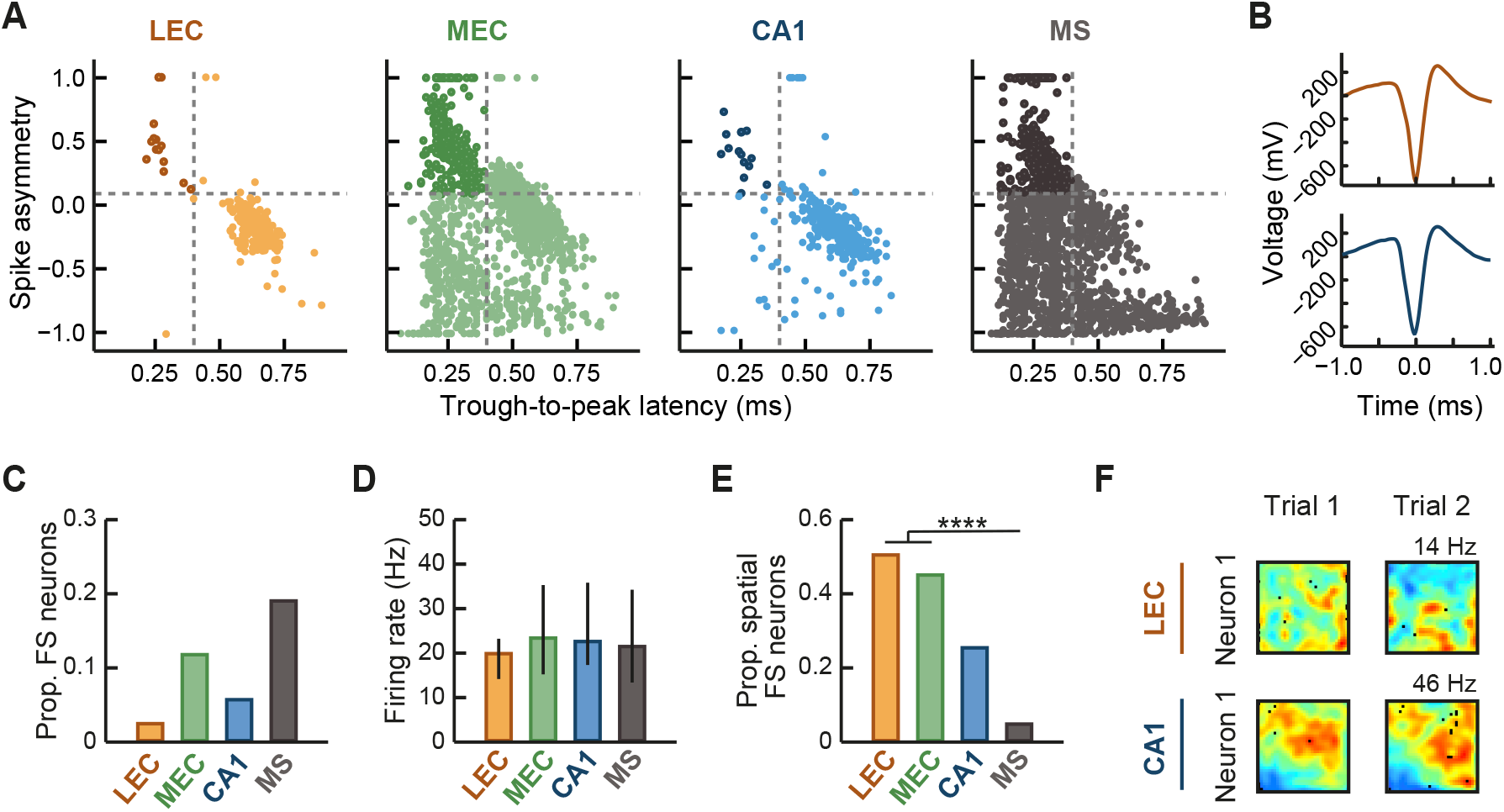
A fraction of fast-spiking neurons in the LEC, MEC and CA1 exhibiting spatial selectivity. **(A)** Trough-to-peak latency and asymmetry of the spike waveform from all recorded neurons in LEC, MEC, CA1 and MS. Fast-spiking neurons (FS) had a spike waveform asymmetry > 0.1, a trough-to-peak latency < 0.4 ms (marked with grey stippled lines) and a mean firing rate > 5 Hz in all open field trials (see Figures S7A and S7B for detail) (n neurons in m mice, LEC: 10 and 5; MEC: 220 and 10; CA1: 13 and 4; MS: 401 and 4). **(B)** Spike waveform of an example FS neuron recorded in LEC (brown) and one recorded in CA1 (blue). **(C)** Proportion of FS neurons recorded in the four areas calculated as the number of FS neurons divided by the number of all active neurons. **(D)** The mean firing rates were similar in FS neurons from the four areas (p > 0.05, H = 6.6563, df = 3, Kruskal-Wallis ANOVA). **(E)** The proportion of spatial FS neurons was similar in LEC and MEC, and higher than that in the MS in both regions (p < 0.0001, X-squared = 153.49, df = 3, χ^2^ test). **(F)** Color-coded rate maps of two example spatial FS neurons, one recorded in the LEC (top) and the other in CA1 (bottom). Maps are scaled from dark blue (silent) to red (peak rate, indicated on top of each map). Pixels not sampled are black. Note that rate maps are stable across two consecutive open field trials in the same box. ****p < 0.0001.

### Local and distal context changes trigger partial remapping in the LEC and CA1

Given that LEC neurons exhibited spatial tuning, we wondered how LEC representations of different spatial contexts compare to those in CA1. Since local and distal cues were shown to exert differential control over neuronal firing in different hippocampal-parahippocampal regions^49–51^, we used two paradigms in which the contribution of distinct local or distal spatial contexts could be assessed. In the “two-box experiment”, recordings were performed in two square boxes (Figures 4A and 4B). Box “a” had black walls and a black floor with a white cue card, box “b” was identical to box a, except that the walls and floor were white and the cue card was black. Henceforth, changes of the box color are referred to as local context changes. The two boxes were placed at the same position in the recording room (Figure 4A). In the “two-room experiment”, two identical black boxes (box “c” and “d”) (Figure 4C) were placed in two recording rooms distinct in size and orientation of the distal cues (similar to^52^). The analysis was confined to active spatially modulated neurons as defined in Figure 2F. In the two-box experiment, many spatially modulated LEC neurons changed their preferred firing location when the box was changed, but remained stable between trials in the same box. Thus, the map stability was high between trials in the same box and low between trials in differently-colored boxes (median map stability in the same box (W1, W2): 0.53 and 0.46; in different boxes (A1, A2): 0.14 and 0.16) (Figure 4B). Remarkably, there was a similar effect in the two-room experiment, i.e., when the distal context was changed (median map stability W1 and W2: 0.51 and 0.50; A1 and A2: 0.18 and 0.15) (Figure 4D). The decrease of map stability was accompanied by a higher mean firing rate change in both the two-box and the two-room experiments (Figures S3A–S3C). LEC remapping was apparent in the dataset from each mouse (Figures S4A and S4B). We divided LEC neurons into three groups: discriminating (high map stability within and low map stability across contexts), stable (high map stability within and across contexts) and unstable (low map stability within and across contexts, see STAR Methods for detail). In both paradigms, the majority of LEC neurons were discriminating, however, the proportion of neurons that distinguished between the two boxes was significantly higher than that of neurons that distinguished between the two rooms (p < 0.0001, X-squared = 207.27, df = 5, χ^2^ test) (Figure 4E, see Figures S4C and S4D for results in two individual mice). Importantly, this effect was not dependent on the threshold used to categorize the neurons (Figure S5A).

**Figure 4.**
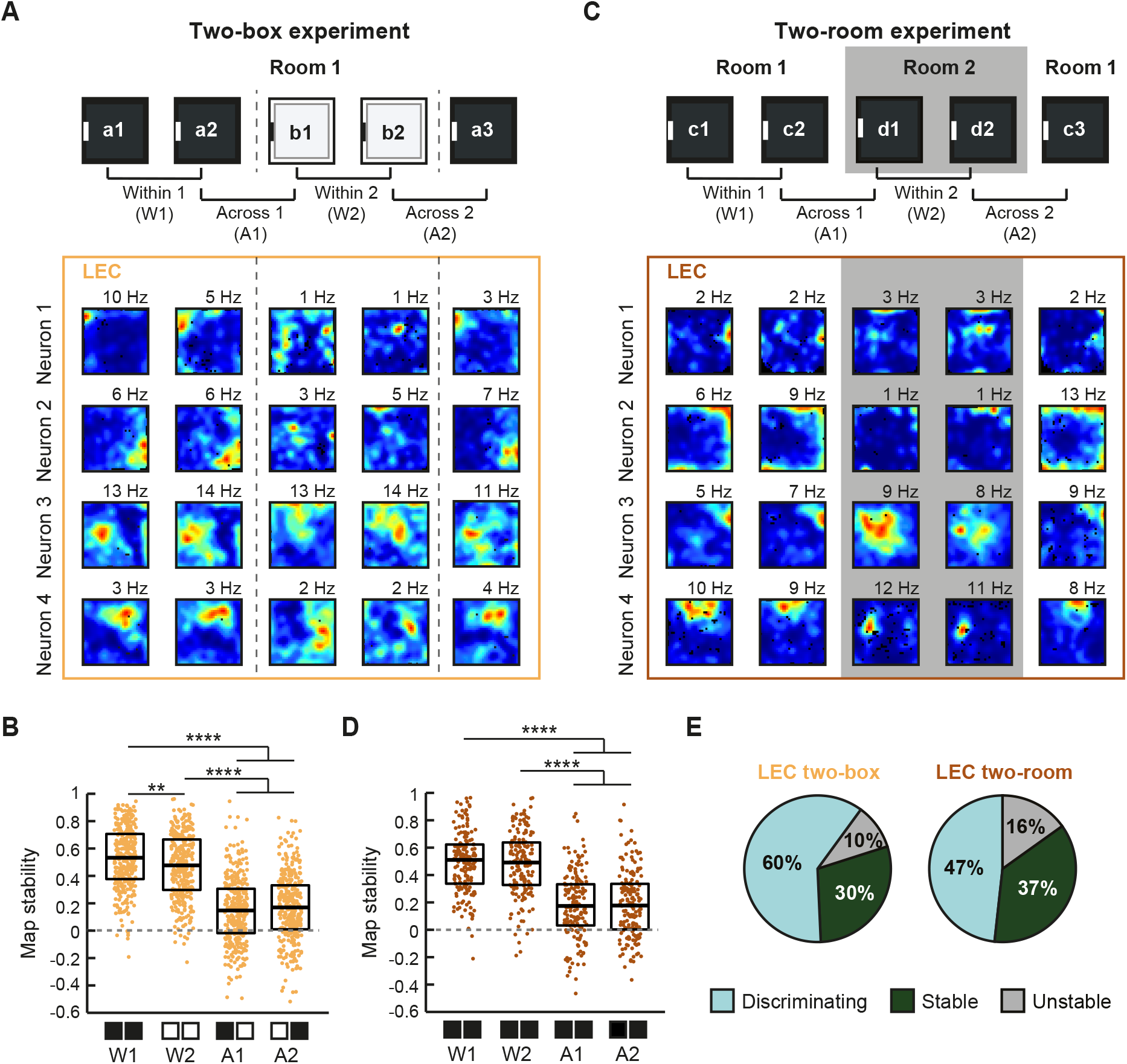
LEC spatially modulated neurons remapping between proximal and distal contexts. **(A)** Color-coded rate maps from representative neurons recorded in the two-box recording setting. (Top) Experimental setting. Recordings were performed by alternating between a black (box a) and white (box b) open field box that were placed in the same position in the room. Recordings were performed in either the depicted protocol or in an “a1-b1-b2-a2-a3” design (see STAR Methods for detail). (Bottom) Color-coded rate maps from four neurons are scaled from dark blue (silent) to red (peak rate, indicated on top of each map). Pixels not sampled are black. Stippled lines delineate the change of boxes. **(B)** The map stability of spatially modulated neurons is higher between trials in the same-coloured box (W1, W2) than between trials in differently-coloured boxes (A1, A2) (p < 0.0001, H = 425.55, df = 3, Kruskal-Wallis ANOVA, n = 307 spatially modulated neurons from 13 mice). **(C)** Same as in **(A)**, but the experiments were carried out in two rooms, as indicated. Two identical black boxes were used. The two black boxes did not induce remapping when placed in the same room (see Figure S6). Recordings were performed in either the depicted protocol or in an “c1-d1-d2-c2-c3” design. **(D)** The map stability of spatially modulated neurons is higher between trials performed in the same room compared to trials performed in different rooms (p < 0.0001, H = 197.29, df = 3, Kruskal-Wallis ANOVA, n = 189 neurons from 5 mice). See Figure S4 for data from two mice recorded in both settings. **(E)** Neurons with high map stability within (> 0.4) and a discrimination index > 0.25 across contexts were categorized as discriminating. The discrimination index was calculated as the difference between the map stability within and across contexts (median(W1,W2)-median(A1,A2)). Neurons were considered “stable” when exhibiting high map stability within and across contexts, but “unstable” when map stability within and across contexts was low (see STAR Methods for detail). Holm-Bonferroni correction was applied for multiple comparisons. Boxplots show medians and interquartile ranges, and each point represents an individual neuron. **p < 0.01, ****p<0.0001.

In line with the reported partial remapping, we observed a decrease in the population vector correlation (two-box: W1, 0.68; A1, 0.30; two-room: W1, 0.67; A1, 0.30) (Figure S5C). We ruled out that the decrease in map stability across contexts was due to a rotation of the map in either paradigm (Figures S5E-S5H). Finally, to ensure that the remapping in the two-box and the two-room experiments was not due to other variables such as olfactory or tactile cues, we performed control recordings identical to those in the two-box experiment, but used two identical black boxes instead of one black and one white box. There was a significantly higher map stability between trials in the different black boxes than in the two-box and the two-room experiments (Figures S6A-S6C).

For comparison, we next determined the response of CA1 place cells in both paradigms. CA1 place cells displayed robust remapping. Specifically, they were more stable between trials in the same context than between trials in different contexts (median map stability, two-box: W1 and W2, 0.76 and 0.72; A1 and A2, 0.15 and 0.15; two-room: W1 and W2, 0.73 and 0.71; A1 and A2, 0.15 and 0.11) (Figures 5A-5D). The map stability difference was accompanied by a mean firing rate change (Figure S3D-S3F). Unlike in the LEC, the proportion of discriminating neurons was similar in the two-box and two-room experiments (Figure 5E, see proportions at different thresholds in Figure S5B). As for the LEC, there was a decreased population vector correlation across contexts compared to trials in the same context in both experiments (two-box: W1, 0.78; A1, 0.26; two-room: W1, 0.73; A1, 0.14) (Figure S5D). Finally, CA1 neurons did not remap between trials in two black boxes in the same room (FigureS S6A, S6D-S6E), and CA1 remapping was not due to a rotation of the map (FigureS S5E-S5H). These experiments reveal, firstly, that under the conditions tested here when there is partial remapping in CA1, there is also partial remapping in LEC, and secondly, that remapping in LEC is triggered by local and distal changes in the environment.

**Figure 5.**
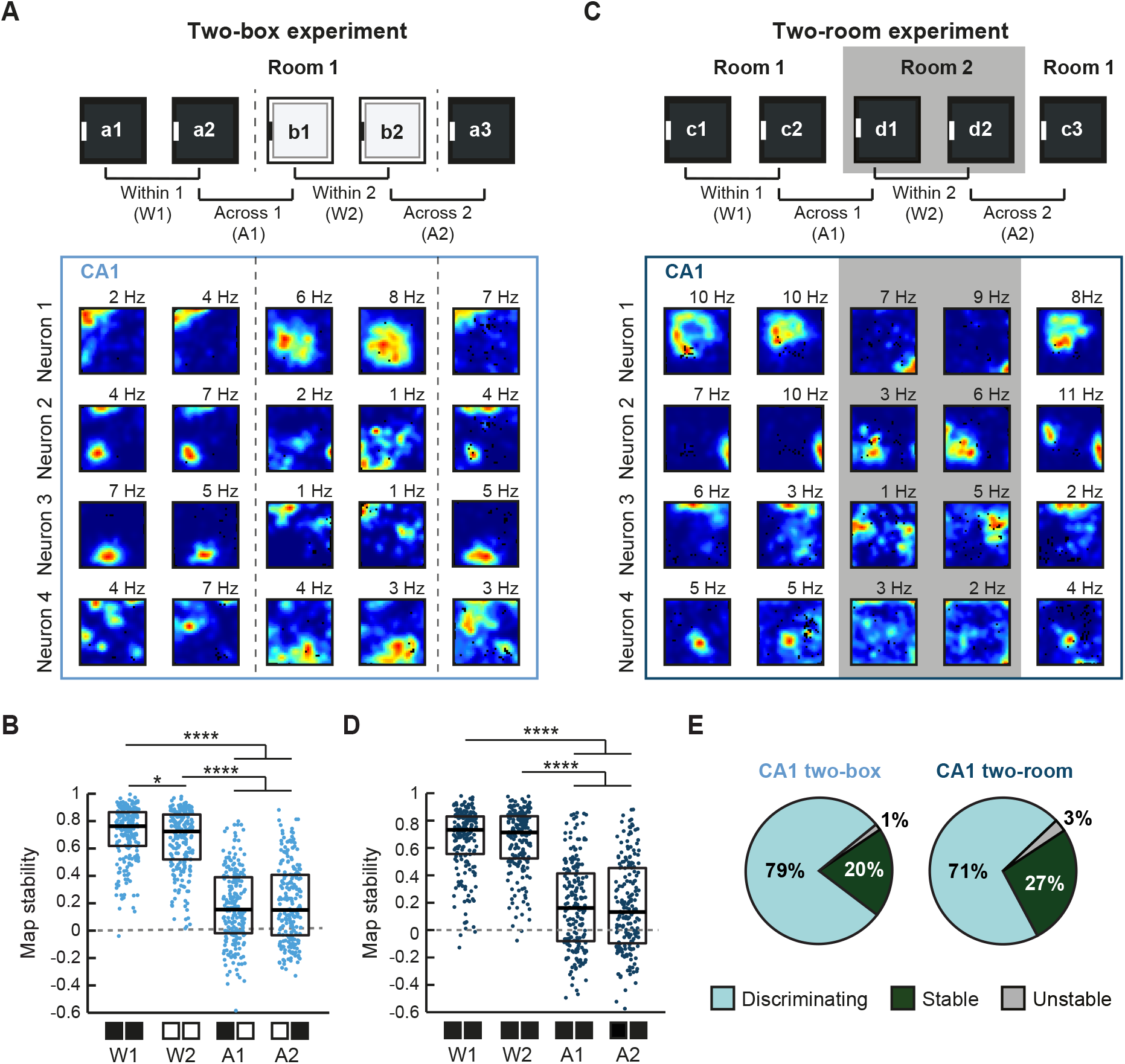
Global remapping in CA1 place cells following a proximal or distal context change. **(A)** Color-coded rate maps from representative neurons recorded in the two-box recording setting. (Top) Experimental setting. Recordings were performed by alternating between a black (box a) and white (box b) open field boxes that were placed in the same position in the room. Recordings were performed in either the depicted protocol or in an “a1-b1-b2-a2-a3” design (see STAR Methods for detail). (Bottom) Color-coded rate maps from four neurons are scaled from dark blue (silent) to red (peak rate, indicated on top of each map). Pixels not sampled are black. Stippled lines delineate the change of boxes. **(B)** The map stability of CA1 place cells is higher between trials in the same-coloured box (W1, W2) than between trials in differently-coloured boxes (A1, A2) (p < 0.0001, H = 446.09, df = 3, Kruskal-Wallis ANOVA, n = 212 place cells from 4 mice). **(C)** Same as in **(A)**, but the experiments were carried out in two rooms, as indicated. Two identical black boxes were used. The two black boxes did not induce remapping when placed in the same room (see Figure S6). Recordings were performed in either the depicted protocol or in an “c1-d1-d2-c2-c3” design. **(D)** The map stability of place cells is higher between trials performed in the same room compared to trials performed in different rooms (p < 0.0001, H = 321.62, df = 3, Kruskal-Wallis ANOVA, n = 181 neurons from 4 mice). **(E)** Discriminating, stable and unstable neurons were categorized as in Figure 4E. Note that a similar proportion of CA1 place cells distinguished between contexts in the two paradigms (p>0.05, X-squared = 505.36, df = 5, χ^2^ test). Holm-Bonferroni correction was applied for multiple comparisons. Boxplots show medians and interquartile ranges, and each point represents an individual neuron. *p < 0.05, ****p < 0.0001.

### Spatially modulated FS neurons in the LEC and CA1 remap upon context changes

As indicated above, FS neurons in LEC and CA1 exhibited spatial selectivity and high stability across trials in the same context. We hence asked whether this stability is maintained across context or whether they remap. Indeed, LEC spatially modulated FS neurons reorganized their firing upon changes in the environment (Figure 6A, see FigureS S7A and S7C for results in the two-room experiment). This reorganization was manifest as a decrease in map stability across contexts, whereas map stability remained high within the same context (median: two-box: W1 and W2, 0.62 and 0.54; A1 and A2, 0.27 and 0.32) (Figure 6B). In the two-room experiment, a similar trend was observed (W1 and W2, 0.62 and 0.60; A1 and A2, 0.47 and 0.37; the low number of neurons (n = 9) does not allow for statistical analysis). In CA1, there was a sharp decrease in map stability of spatial FS neurons between trials in different contexts in both paradigms (median: two-box: W1 and W2, 0.60 and 0.59, A1 and A2, 0.14 and 0.12; two-room: W1 and W2, 0.60 and 0.62; A1 and A2, 0.15 and 0.13) (Figures 6C, 6D, S7B, S7D and S7E).

**Figure 6.**
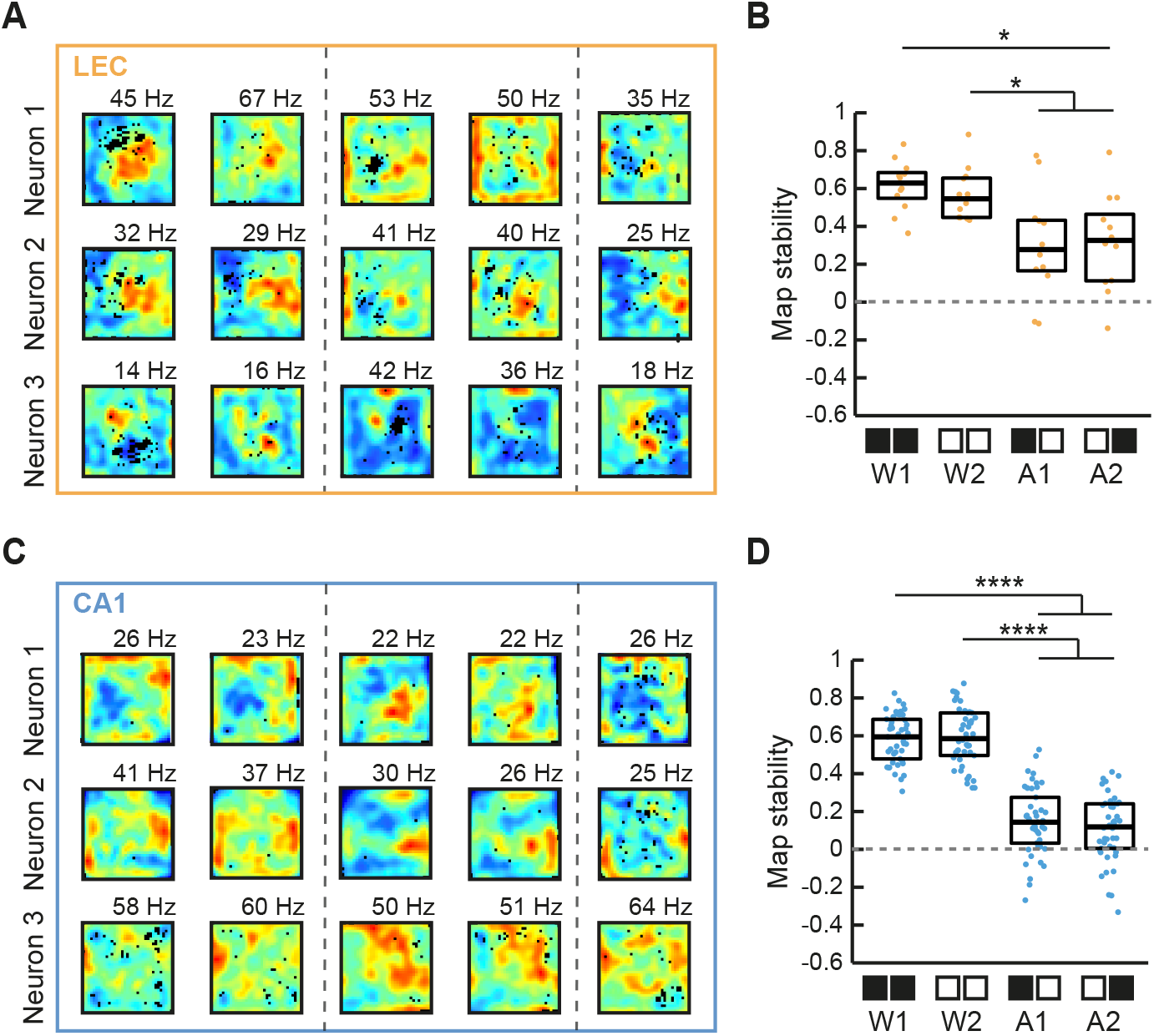
Remapping of spatially modulated FS neurons in LEC and CA1 upon context changes. **(A)** Color-coded rate maps from discriminating LEC FS neurons recorded in the two-box experiment (see Figure 4A for detail). Maps are scaled from dark blue (silent) to red (peak rate, indicated on top of each map). Pixels not sampled are black. Stippled lines delineate the change of boxes. **(B)** The map stability of LEC FS spatially modulated neurons is higher between trials in the same box (W1, W2) than between different boxes (A1, A2) (p < 0.05, H = 16.674, df = 3, Kruskal-Wallis ANOVA, n = 12 neurons from 5 mice). **(C)** Same as **(A)** from three discriminating CA1 FS neurons. **(D)** The map stability of CA1 FS spatially modulated neurons is higher between trials in the same box than between trials in different boxes (p < 0.001, H = 125.99, df = 3, Kruskal-Wallis ANOVA, n = 45 neurons from 4 mice). Holm-Bonferroni correction procedure was applied for multiple comparisons. Boxplots show medians and interquartile ranges, and each point represents an individual neuron. *p < 0.05, ****p < 0.0001.

### Context information is processed independently of object information in the LEC

Behavioral studies following lesions or neuronal inactivation showed that the LEC is involved in object-context and object-place-context associations^29–31^. However, how the LEC encodes object and context information in spontaneous exploration conditions is still an open question. Thus, we recorded from 6 LEC-implanted mice and 3 CA1-implanted mice while they foraged for food in an open field with one object present (Figure 7A). The object and/or local context configurations were modified such to allow for the following comparisons between consecutive trials: (1) same context (i.e., the same box) and different object location (sCdO); (2) different context (i.e., black and white box) and same object location (dCsO); (3) same context and same object location (sCsO); and (4) different context and different object location (dCdO).

**Figure 7.**
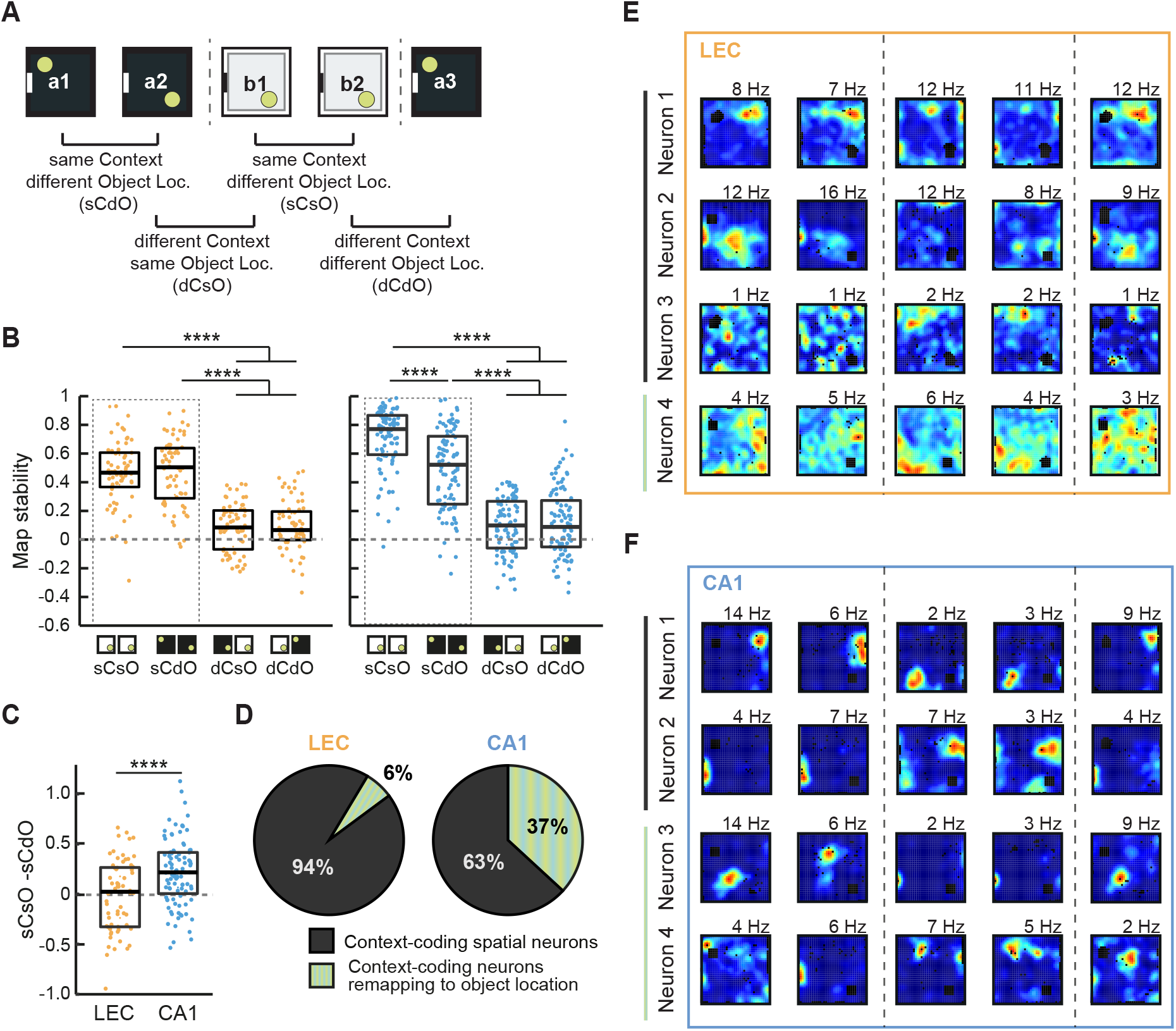
Context-coding neurons remapping to the object location in CA1, but not in the LEC. **(A)** Experimental setting. Recordings were performed by alternating between a black (box a) and a white (box b) open field box that was placed in the same position in the room. Inside the box, an object (green circle) was placed in the indicated position. The experimental setting allowed to compare the firing of LEC neurons between trials in which the context was the same but the object location changed (sCdO), the context changed but the object was in the same location in the box (dCsO), both the context and the object location changed (dCdO) or both the context and the object location were constant (sCsO). **(B)** The map stability of context-coding LEC neurons (left) is higher between trials in the same box (sCsO, sCdO) than between trials in different boxes (dCsO, dCdO) and does not decrease when only the object is moved (sCdO) (p < 0.0001 and p > 0.05, respectively, H = 122.06, df = 3, Kruskal-Wallis ANOVA, n = 64 neurons from 6 mice). In context-coding CA1 neurons (right), the map stability decreases when the object is moved in the same box and when the box is changed (p < 0.0001, H = 206.54, df = 3, Kruskal-Wallis ANOVA, n = 95 neurons from 3 mice). **(C)** The difference in map stability when comparing sCsO trials and sCdO trials is greater in context-coding CA1 neurons than in context-coding LEC neurons (p < 0.0001, W = 8726, Wilcoxon sum rank test). (D) Proportion of context-coding neurons that remap to the object location in LEC (left) and in CA1 (right). ‘Remapping to the object location’ was considered when the decrease in map stability between sCdO trials was greater than 25% of the map stability between sCsO trials. **(E-F)** Color-coded rate maps of LEC neurons (E) or CA1 neurons **(F)** corresponding to each of the categories described in **(D)**. Maps are scaled from blue (silent) to red (peak rate, indicated on top of each map). Pixels not sampled are black. Stippled lines delineate change of boxes. Holm-Bonferroni correction procedure was applied for multiple comparisons. Boxplots show medians and interquartile ranges, and each point represents an individual neuron. **** p < 0.0001.

We first asked whether spatially modulated neurons that are modulated by changes in the local context (i.e., context-coding neurons, 55% of spatially modulated neurons in LEC and 61% in CA1) are modulated additionally by object-related modifications in the environment, i.e., induced by the object location change. Notably, this was the case for CA1 but not for LEC. Specifically, CA1 neurons that remapped to the change of the box color (indicated by a decreased map stability) also remapped to the change of object location (median map stability: sCsO, 0.77; sCdO; 0.52; dCsO, 0.10; dCdO, 0.09) (Figure 7B). In contrast, LEC neurons that remapped to the change of the box color did not exhibit remapping upon changing the object location (median map stability: sCsO, 0.47; sCdO, 0.50; dCsO, 0.09; dCdO, 0.07) (Figure 7B). In accordance, the decrease in map stability between the sCsO and sCdO conditions was much higher in CA1 neurons compared to LEC neurons (median difference: LEC, 0.02; CA1, 0.22) (Figure 7C). Likewise, the proportion of neurons that remapped to the object location change in addition to the context change was higher in CA1 (37%) than in LEC (6%) (Figures 7D and 7E). Together, the results highlight that spatial representations in the LEC are less prone to be altered by other variables (here, object location) in the environment than representations in CA1.

### A heterogeneous population of object cells in the LEC comprises a large fraction of context-independent object identity cells

We next focused on object cells, which we divided into two groups: context-independent object cells (z-score > 2 around the object location in the first three trials) and contextdependent object cells (z-score > 2 around the object location in all black box trials but not in white box trials) (Figures 8A and 8B). In the LEC, object coding in general was sparse. Yet, we were able to identify both context-independent object cells (5.2%) and context-dependent object cells (0.7%). In CA1, object cells also comprised both cell types (context-independent: 5.9%; context-dependent: 2.9%) (Figure 8B). These results indicate that, in an unbiased dataset (tetrodes were turned after each recoding session and the experimenter was blind to the result), there was sparse object coding in the LEC.

**Figure 8.**
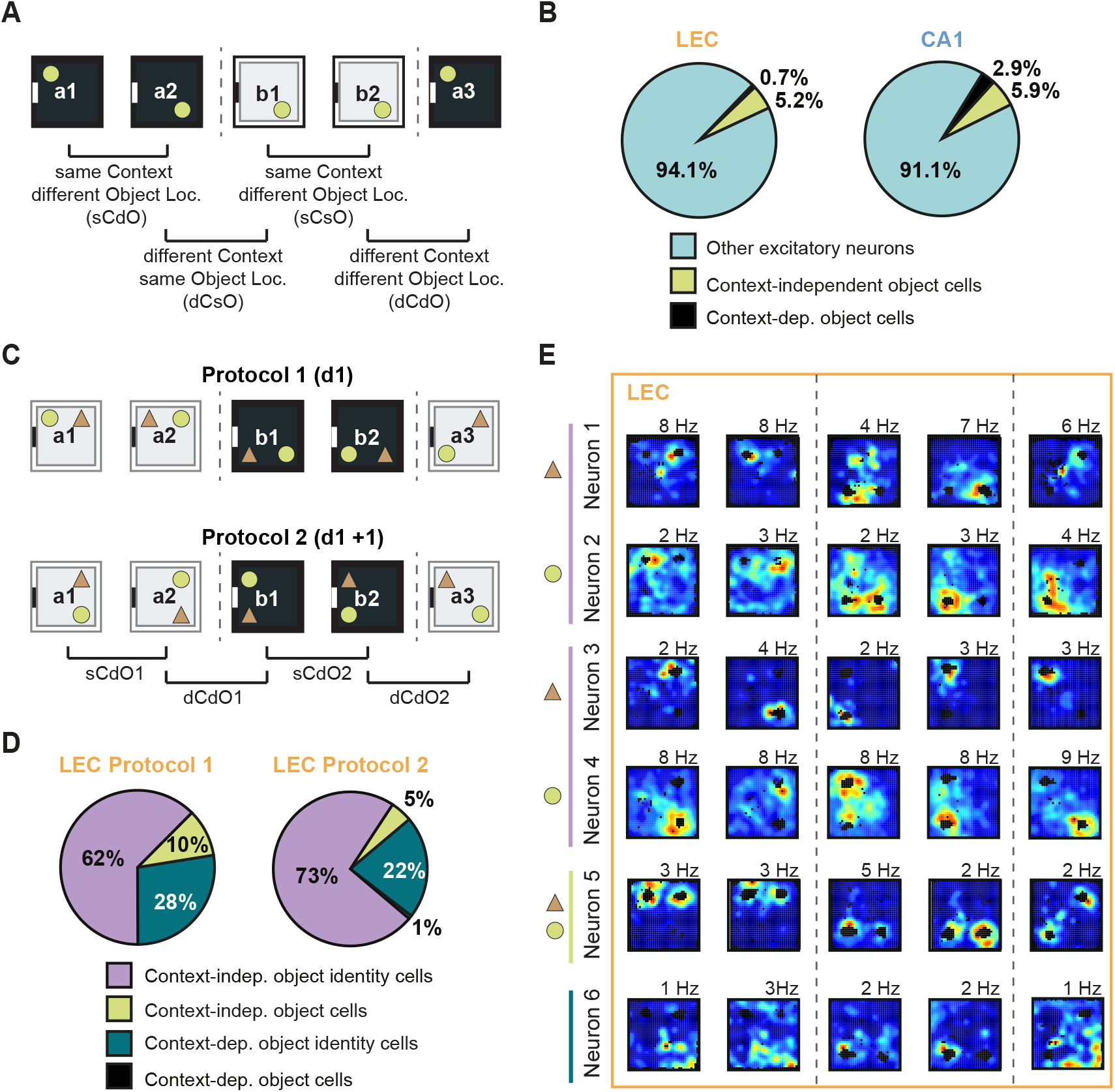
Context-dependent and context-independent encoding for objects and object identities in LEC neurons. **(A)** Experimental setting (see details in Figure 7A). Recordings were performed by alternating between a black (box a) and a white (box b) open field box that was placed in the same position in the room. Inside the box, an object (green circle) was placed in the indicated position. **(B)** Proportion of excitatory LEC (left) and CA1 (right) neurons that are context-independent object cells (green, z-score > 2 in the first three open field trials), contextdependent object cells (black, z-score > 2 only in black box trials) and other excitatory neurons (turquoise) (LEC: 288 neurons in 6 mice, CA1: 204 neurons in 3 mice). **(C)** Experimental settings. In this paradigm, two objects (green circle and brown triangle) were placed at the indicated positions inside the boxes, allowing us to assess object identity coding in different contexts. Protocols 1 and 2 were alternated on consecutive days. The experimental settings allowed to compare the firing of LEC neurons between trials where the context was the same but the location of the objects changed (sCdO1 and sCdO2), and between trials where both the context and the location of the objects were changed (dCdO1 and dCdO2). **(D)** Proportion of all object cells (protocol 1: n = 80 LEC neurons in 3 mice; protocol 2: n = 130 LEC neurons in 3 mice) that are context-dependent object cells (black, z-score > 2 at both object locations only in one box), context-dependent object identity cells (blue, z-score > 2 at one of the object locations only in one box), context-independent object identity cells (violet, z-score > 2 at the location where either the circle or the triangle object are in a2 and a3 trials) and context-independent object cells (green, z-score at both of the object locations > 2 in a2 and a3 open field trials). **(E)** Color-coded rate maps from LEC neurons that are context-independent object identity cells with selectivity for each of the objects in protocols 1 (neurons 1 and 2) and protocol 2 (neurons 3 and 4), one context-independent object cell (neuron 5) and one contextdependent object cell (neuron 6).

As previous studies clearly demonstrated that the LEC supports object coding, object recognition and object-place-context recognition^13,19,20,29–31^, we next actively searched for object cells to further characterize this population. To this end, we recorded preferentially object cells in the LEC of 3 mice employing an experimental setting that allowed us to discern putative effects related to different contexts and object identities (Figure 8C). In this dataset with a higher number of object cells (Figure 8D), we also identified both context-independent and context-dependent object cells. The former constituted the largest fraction, and most importantly, comprised a heterogeneous population. The vast majority of context-independent object cells were selective for one specific object (i.e. object identity cells, e.g. Neurons 1-4 in Figure 8E; 62% and 73% of all object cells in protocols 1 and 2, respectively) whereas a small fraction encoded for both objects, irrespective of their identity (i.e. object cells, e.g. Neuron 5 in Figure 8E, 10% and 5% of all object cells) (see STAR Methods for classification criteria). We also identified context-dependent object cells (e.g. Neuron 6 in Figure 8E). Interestingly, most context-dependent object cells were selective for one specific object (28% and 22%), and virtually no context-dependent neurons were object cells (0% and 1%). Hence, object coding in the LEC is supported by a heterogeneous population of neurons that encode objects both in a context-dependent and in a context-independent fashion.

## Discussion

In this study, we characterized the spatial coding properties of LEC neurons, and explored a putative co-existence and interdependence of object and spatial context coding in this region. We found that excitatory neurons in the anterior/intermediate LEC, but less so in the posterior LEC, were spatially modulated. Spatially modulated LEC neurons exhibited partial remapping upon changes in the local or the distal context. A fraction of FS neurons in the LEC, MEC and CA1 area were also spatially modulated and remapped between different contexts. A series of experimental settings allowed us to discriminate between context-coding, object-coding and object-identity-coding neurons in the LEC. Notably, space and object coding was largely independent of each other. However, we also identified a fraction of conjunctive contextdependent object cells.

Spatial firing of neurons in the LEC was less evident in previous studies in standard open field settings, but based on the published histological data it can be inferred that the recording sites were mostly in the posterior LEC^19,20,22–25,28,53^. Spatial selectivity gradients have been reported also for the hippocampus and the MEC^35,54–56^, and may reflect both a gradient of gene expression and local cellular connectivity as well as different inputs from upstream regions. At this point it can only be speculated as to whether the spatial selectivity gradient can be accounted for, at least in part, by differential inputs in LEC. For instance, there is a pronounced antero-posterior gradient of the perirhinal cortex projections to the LEC^11,26^. Furthermore, postrhinal cortex projections target preferentially more anterior levels of LEC^26^.

A surprising finding in this study is the high proportion of spatially modulated neurons in an experimental setting involving spontaneous exploration in a standard empty open field. LEC neurons that exhibit spatial firing to some degree have been reported before^19,20,27^. However, these neurons with “place-like firing” as termed by the authors, were found in a setting with objects rather than an empty open field. The previous studies already implicated that there is not a clear-cut division regarding spatial versus non-spatial representation in the MEC and LEC, respectively^39,40,57^. This prompts the pressing question to which extent spatially modulated neurons in MEC and LEC contribute to place field firing in the hippocampus. Lesion studies of either MEC or LEC did not abolish completely the spatial tuning in the hippocampus^53,58^. It is tempting to speculate that following MEC lesions, the remaining spatial input from the LEC could maintain spatial coding in CA1. In agreement with this possibility, it was reported in a recent computational study that input from weakly spatially modulated neurons in the entorhinal cortex sufficed to trigger place field formation in hippocampal place cells^59^. Of course, a completely different scenario can be envisaged, as hippocampal neurons also provide input to the deep layers of LEC. Thus, the spatial firing of LEC neurons could also be inherited from CA1/subiculum.

Given that we found spatially modulated neurons in the LEC, we wondered whether and, if so, how the LEC map may incorporate changes in the spatial configuration of the environment. It has been repeatedly shown that environmental changes trigger partial or global remapping in the hippocampus^49,52^. A major conclusion of our study is that a change of the spatial context (be it local or distal) triggers partial remapping in the LEC. How do these results compare to previous studies? For instance, Lu and colleagues reported that the change of the wall color of the open field box triggered rate remapping in the hippocampus, but not in the LEC^53^. In the setting of that study, hippocampal spatial representations were stable, whilst in our study, the change of both the wall and floor color triggered partial remapping in the hippocampus. We found that in these conditions, upstream of the hippocampus in the LEC, there is also a change in the spatial representation. In yet another study, the authors recorded LEC neuronal activity when the environmental changes consisted in a mismatch between the “proximal” and “distal” cues of the environment (i.e., proximal cues were rotated counterclockwise and distal cues were rotated clockwise)^60^. Notably, the recorded neurons were predominantly non-spatial, a small fraction (9%) was modulated by proximal cues and virtually no neurons were modulated by distal cues. As our recording location within the LEC allowed us to record from a much larger proportion of spatially modulated neurons, we asked how a complete change of the local or distal spatial context is incorporated into the LEC spatial map. Thus, we found that a high proportion of spatial LEC neurons remap upon changes in both the distal (47%) or the local context (60%), providing evidence that both spatial frameworks are represented in the LEC.

Spatial modulation of LEC neuronal activity was not restricted to excitatory neurons, but extended to FS neurons. It has often been considered that FS neurons subserve input-gain control in neurons in different brain areas (including hippocampus, MEC, visual cortex, etc.)^44–46^. Thus, in contrast to excitatory neurons that encoded environment-related features, FS neurons were significantly less tuned. In this study we found that a sizeable fraction of FS neurons in MEC, hippocampus and LEC, but not in the MS, were spatially modulated in our experimental setting. This indicates that, in the hippocampal-entorhinal regions, FS neurons are involved in spatial coding supporting a scenario according to which there is dedicated ensemble-specific inhibition. In previous studies, it became apparent that FS neurons may encode specific features, particularly in more complex environmental settings, e.g., upon introduction of objects or rewards^47,61^. Here, we demonstrated that spatial FS neurons in LEC and CA1 remapped in different spatial contexts. This further substantiates the notion that FS neurons contribute to feature coding in the networks that they are embedded in^18,43,44,47,62^.

Previous behavioral studies demonstrated that the LEC is crucial for object-context associative memory^29–31^. These studies raise the important question whether the association of object and context-related information is carried out in the LEC or whether the separately transmitted information is associated downstream in the hippocampus. We explored how object and context representations are generated in the LEC at the single-neuron level. Our results demonstrate that, during random foraging, spatial coding of different contexts is independent from object coding in the LEC. Furthermore, we identified a variety of neurons encoding specific objects that was by and large context-independent. The existence of context-independent non-spatial codes upstream of the hippocampus confirms predictions postulated by the cognitive map theory^36,37^. In addition, we identified a small population of LEC neurons that supported context-dependent object coding. This indicates that some integration of the object and context information is carried out in the LEC. At this point one can only speculate about the identity of the source neurons that provide the information to these neurons that enable conjunctive coding. Possible scenarios include: (1) the input is provided both by perirhinal and postrhinal projections onto the same LEC neurons^26^, (2) back-projections from the hippocampal formation convey the joint object and context information (3) distinct object and context coding neurons in the LEC provide local input to conjunctive context-dependent object neurons (4) projections from MEC convey spatial information onto object-coding LEC neurons^63,64^.

Finally, the identification of context-dependent object cells during spontaneous exploration prompts future inquiries as to whether conjunctive coding becomes more prominent in LEC-dependent memory tasks. For instance, one could envisage a scenario in which the recruitment of separate and conjunctive codes in the LEC is flexible and task-dependent. This may hold true also for the hippocampus that, akin to the LEC, harbors neurons that perform separate and conjunctive object and context coding. Thus, a rigid allocation of spatial versus non-spatial processing to a particular area may be overly simplistic.

## Supporting information

Key resources table

Statistics and quantification tables

## Acknowledgements

The authors would like to thank Dr. Antonio Caputi and Dr. Kevin Allen for input and discussion on previous versions of the manuscript, and Sofiya Zbranska, Miruna Rascu and Tahir Mehmood for technical assistance. This work was funded by the DFG grant MO 432/17-1, by the BWST_ISF 2018-036 grant, by the GIF_I-151-421.10-2017 German-Israeli Grant, and by the ERA-NET Neuron NMDA-PSY to H.M., the Baden-Wurttemberg Stiftung elite program for postdocs, the Olympia Morata program, and a Marie Sklodowska Curie individual fellowship (H2020-MSCA-IF-708295) and DFG Emmy Noether Programme grant to M.I.S.; the China Scholarship Council (CSC) doctoral fellowship to X.H. and a Boehringer Ingelheim Fonds PhD fellowship to I.B-O.

## Author contribution statement

Conceptualisation: the project was developed and discussed throughout the entire period of execution by X.H., M.I.S, I.B-O. and H.M.; LEC surgeries were established by M.I.S and performed by X.H; LEC and CA1 recordings: X.H. and I.B-O.; Surgeries and recordings for MS experiments were performed by D.A.A.M., M.I.S. and N.B. Data analysis was performed by I.B-O., X.H. and M.I.S.; Decoding analysis: C.L.; Supervision and coordination of the project: M.I.S. and H.M.; Writing: first draft was generated by I.B-O., M.I.S. and H.M.; generation of the manuscript was performed by I.B-O., X.H., M.I.S. and H.M.; generation of the figures was performed by I.B-O.; manuscript editing and proofreading was performed by all authors.

## STAR Methods

### RESOURCE AVAILABILITY

#### Lead contact

Further information and requests for resources and reagents should be directed to and will be fulfilled by the lead contact, Prof. Dr. Hannah Monyer.

#### Materials availability

This study did not generate new unique reagents.

#### Data and code availability

The datasets generated or re-analyzed during the current study are available from the corresponding author upon request. All original code from this study are available from the corresponding author upon request. Any additional information required to re-analyze the data reported in this paper is available from the lead contact upon request.

### EXPERIMENTAL MODEL AND SUBJECT DETAILS

LEC and CA1 recordings were performed in naïve wild-type male mice maintained on a C57BL/6N background (n = 33). For the optogenetic identification of medial septal neurons, we used naïve male PV^Cre^ mice (n = 4 mice) maintained on a C57BL/6N background^65^. Mice were 14-18 weeks old at the start of the experiments. Mice were kept single-housed on a 12-hour dark/light cycle with water *ad libitum*. Training and recordings were performed during the light phase of the cycle. At the start of the electrophysiological experiments, mice were food restricted and maintained at >85% of their free-food body weight. All experiments were approved by the Regierungspräsidium Karlsruhe and were in compliance with the European guidelines for the care and use of laboratory animals. CA1 and MEC recording data in Figures 2 and 3 were reused from ref.^66^ (CA1 recording data of control animals) and ref.^67^ (MEC recording data).

### METHOD DETAILS

#### Microdrive assembly and implantation

Prior to implantation, tetrodes were constructed by twisting tungsten wire (California Fine Wire Company). They were subsequently mounted into custom-designed microdrives (Axona) that allowed the individual movement of the tetrodes. For the experiments that examined the antero-posterior gradient of spatial selectivity in LEC (Figure 1), we used 3-tetrode microdrives and implanted each of the 3 tetrodes at the following antero-posterior (AP) coordinates from bregma: anterior tetrode: −2.92 mm; intermediate tetrode: −3.80 mm; posterior tetrode: −4.36 mm. All 3 tetrodes were implanted at ML −4.3 mm and DV −2.3 mm. In the rest of the experiments, implantations in the MS and LEC were performed using 4-tetrode microdrives, and implantations in CA1 were performed using either 4- or 12-tetrode microdrives. For one mouse from experiments shown in Figure 2 we used a 1-tetrode microdrive. For the MS recordings, an optic fiber (125-0.22_17mm_ZF1.25-FLT, Doric lenses) was additionally mounted into the microdrives for optogenetic stimulation and subsequent detection of parvalbumin-positive (PV^+^) neurons. During surgery, mice were mounted in a stereotaxic apparatus and kept under isoflurane anesthesia (1.0%–2.5%). A craniotomy was made above the site(s) of implantation and the tetrodes were subsequently implanted. For LEC recordings, tetrodes were implanted at AP: −3.80 mm, ML: −4.30 mm, DV: −1.8 mm with a 4° angle in the lateral direction. For CA1 recordings, tetrodes were implanted bilaterally at AP: −1.80 mm, ML: ±1.7 mm and DV: −0.90 mm. A subset of mice (2 mice from Figure 5) was implanted unilaterally in the hippocampus at ML: ±1.7. For MS recordings, prior to the microdrive implantation, a total of 400nL of pAAV-EF1a-double floxed-hChR2(H134R)-mCherry-WPRE-HGHpA was injected from a glass pipette into the MS. The first 200 nL were injected at AP: +1.00 mm, ML: ±0.00 mm, DV: − 3.80 and after 4 min an additional 200 nL were injected at AP: +1.00 mm, ML: ±0.00 mm, DV: − 4.10. After another 4 min the glass pipette was removed. The virus was a gift from Karl Deisseroth and obtained from Addgene (Cat# 20297). The tetrodes and optic fiber were implanted at AP: +1.00 mm, ML: −0.30 mm, DV −3.8 mm and an angle of 3° towards the midline. Mice recovered for at least one week prior to the start of the food restriction and experiments.

#### In vivo electrophysiological recordings: data acquisition equipment

Electrophysiological signals were recorded and digitized with a RHD2000 (Intan Technologies) data acquisition system, using a sampling frequency of 20 kHz. Microdrives implanted on the head of the mice were connected to the data acquisition system by an ultrathin SPI cable (Intan Technologies) and the signal was amplified with an RHD head stage (Intan Technologies). The position of the mouse in the open field arena was triangulated from the position of three LEDs (red, green, blue) that were connected to the head stage. The position of each LED was recorded with a camera (DFK 23GM021, The Imaging Source) using the ’positrack’ software (https://github.com/kevin-allen/positrack) at a sampling frequency of 50 Hz. Tracking signal was connected to the data acquisition system and sampling time stamps of each position were recorded along with the electrophysiological signal using the ‘ktan’ software (https://github.com/kevin-allen/ktan). In addition, for MS recordings where laser stimulation was performed, time stamps of each laser pulse were recorded along with the electrophysiological signal.

#### In vivo electrophysiological recordings: experimental design and recording chambers

At least one week after surgery, all mice were food restricted and trained to forage for sugar pellets (Ain-76A Rodent Tablet 5 mg, TestDiet™) randomly distributed by a dispenser (Med Associates) in a black open field arena (70 cm x 70 cm x 30 cm) with a polarizing white cue card (21 cm x 30 cm) situated in the center of the left side in relation to the setup’s opening. Mice were first trained in two or three 20-minute open field trials per day for at least 3 days without connecting their implanted microdrive to the data acquisition system, until they learned to eat most of the sugar pellets delivered. Before, after and in-between trials, mice were transferred to a rest box (35 cm x 35 cm) for 10 minutes. During training and in all the recording protocols, the open field arena was cleaned with water while the mouse was in the rest box, to avoid any potential olfactory cues. Subsequently, mice were habituated to forage for sugar pellets with their implanted microdrive connected to the data acquisition system via the amplifier and cable in two or three 20-minute open field trials per day for at least 3 days. Tetrodes were lowered gradually during training until clear action potentials (spikes) were observed in the local field potential. When mice showed good coverage in all the open field trials for at least two consecutive training days, the corresponding recording protocol started.

##### Assessment of spatial firing in the LEC (Figures 1–3)

For experiments assessing the antero-posterior gradient of spatial selectivity (n = 5 mice) and the proportion of spatially modulated neurons in each of the hippocampal-parahippocampal regions (n = 5 mice implanted in LEC, n = 10 mice implanted in MEC, n = 4 mice implanted in CA1, n = 4 mice implanted in MS), recordings were performed during random foraging in two 20-minute trials in a black square box (70 cm x 70 cm x 30 cm), separated by a 10-minute period in a rest box. After every recording day with good coverage and with intact electrophysiological and tracking data, tetrodes were lowered by about 50 μm to obtain new neurons for the next recording session. This procedure was repeated until no more neurons could be recorded and/or the electrophysiological signal indicated that the tetrodes went past the region of interest. To ensure that the neurons recorded in the LEC corresponded to the superficial layers (Figures 2 and 3), we only considered mice in which tetrodes tips were in superficial LEC layers and only included the last 5 recording sessions before the end of the experiment (perfusion). Mice with tetrode tips located in the deep layers of LEC were excluded from the analysis (n = 3 mice). In order to identify the location of the MS and therefore avoid recording from the lateral septum (as neurons in the lateral septum have some spatial coding properties^40^), for the MS recordings, optogenetic stimulation was performed to tag the PV^+^ neurons located within the MS (similar to the methodology we previously used in^43^ to tag MEC PV^+^ neurons). During one of the open field trials (sham and stim conditions were counterbalanced), a 5-ms 473 nm wavelength laser pulse was delivered at a frequency of ~6.67 Hz (intervals of 145 – 155 ms) via a patch cord (100 mm multimode fiber core, Doric Lenses) connected to the optic fiber within the microdrive. A neuron was considered tagged if its firing rate rose to above 4 SD of its baseline firing (50 ms interval immediately prior to laser onset) within 5 ms of laser onset. Analysis was performed on the sham conditions, however, only tagged neurons or neurons co-recorded on the same tetrode and in the same recording session as tagged PV^+^ (and hence likely corresponding to the most medial part of the MS) were included in the analysis (n = 4 mice).

##### Assessment of context-dependent spatial firing of LEC neurons

As for the previous set of experiments, mice with tetrodes implanted into LEC (n = 5) or CA1 (n = 4) were trained in a black square box (70 cm x 70 cm x 30 cm) where sugar pellets were randomly dispensed by an automatic pellet feeder (Med Associates). The training protocol was as follows: 3 days of training unplugged from the data acquisition system in three 20-minute open field trials; at least 3 days of training connected to the data acquisition system in three 20-minute open field trials; at least 5 days in five 20-minute trials. In-between trials and before and after a daily trial series, mice were subjected to 10-minute periods in a rest box. Recordings began when mice showed good coverage of the recording arena in each of the five trials and spikes from at least 5 neurons could be isolated. The recording protocol consisted of three phases, in the following order: one day of “two-black box experiment” (control experiment); seven days of “two-box experiment” and at least seven days of “two-room experiment” (see detailed protocol below).

###### Two-black box experiment (Figure S6)

Mice were firstly subjected to a control experiment with two trials in one black box (70 cm x 70 cm x 30 cm, the same one that was used for training, box A), then two trials in a second identical black box (box B) and one last trial in box A. The two black boxes were placed in the same location within the recording room and oriented such that the white cue card was in the same position. In-between trials and before and after a daily trial series, mice were subjected to 10-minute periods in a rest box. This allowed us to reliably track the action potentials of neurons that strongly reduced their firing rate in one of the two environments (see e.g. refs.^50,68,69^).

###### Two-box experiment (Figures 4–6)

Mice underwent 7 days of recordings in the two-box experiment (see Figures 4A and 5A for schematic). During the first 5 days, we employed an “A A B B A” design: mice foraged for sugar pellets in two 20-minute trials in a black square box (70 cm x 70 cm x 30 cm) separated by 10-minute rest box periods. Subsequently, the black box was substituted by a white square box (70 cm x 70 cm x 30 cm) with a black cue card situated at the same location as the white cue card was in the black box (i.e., on the left side in relation to the opening of the setup). Recordings in the white box were performed for two 20-minute trials. The fifth 20-min trial was performed in the black box. The white box was novel to the mouse at the start of the experiment. The boxes were cleaned with water in-between trials. The cable was not detached from the mouse throughout the daily recording series. A rest box period was included before the first and after the last open field trial, to reliably track neuronal activity throughout the daily recording series. On days 6 and 7 of the two-box experiment, recordings were performed as described above but in an “A B B A A” design. This was to exclude the possibility that the observed effects were driven by the order of trials or changes over time. A subset of LEC-implanted mice (n = 9 out of 13 mice) only underwent the general training and this protocol (without the two-black box experiment and the two-room experiment).

###### Two-room experiment (Figures 4–6)

Recordings were subsequently performed in the two-room experiment for 7 days (illustrated schematically in Figures 4C and 5C) in a procedure identical to that in the two-box experiment, except that two identical black boxes (70 cm x 70 cm x 30 cm) were placed in two different rooms. As above, we recorded for 5 days in an “A A B B A” design followed by 2 days in an “A B B A A” design. One of the rooms was novel to the mouse at the start of the experiment, the other was introduced to the mouse during previous training and the two-box experiment. The rest box placed on a cart was used as a vehicle to transport the mouse between rooms. The recording cable was not detached from the mouse during the daily recording series, and manually stabilized by the experimenter during the transport. After the transport, a 10-minute rest period started. In the two rooms, the identical black boxes had a white cue card that was situated at the same angle and direction in relation to the opening of the setup.

After 7 days of recording in this standard protocol, recordings continued using the “A A B B A” and the “A B B A A” design on alternating days for 1-15 days, until tetrodes passed layer I and no action potentials could be detected. Tetrodes were gradually lowered by ~50 μm after each daily recording session with good coverage in all five trials, and only sessions with good coverage were included into the data analysis.

##### Object in context experiment (Figure 7)

A separate set of mice with microdrives implanted into LEC (n = 6 mice) or CA1 (n = 3 mice) was subjected to the general training as in previous experiments, with the difference that a cylindrical object (8.5 cm in diameter, 15 cm in height) was placed in the open field during the last 5 days of training. During a daily session, the object was either located in the bottom left corner (trials 1 and 5) or the top right corner (trials 2-4) at a distance of 20 cm from the respective corner.

The object and the open field were cleaned with water after each trial. Open field trials were separated by 10-minute rest box periods, and a rest box period was included before the first and after the last open field trials. After mice showed good coverage in all trials and spikes from at least 5 neurons could be isolated, recordings in the object in context experiment started. Specifically, recordings were performed following the same recording protocol as in the two-box experiment, but with a cylindrical object inside of the open field boxes (see Figure 7A for schematic). During the first 5 days the black box (box A) and the white box (box B) were arranged in an “A A B B A” design and the object location varied between the bottom left (bl) and the top right (tr) corner in the following sequence: “bl tr tr tr bl”. The white box was novel to the mouse at the start of the experiment. On days 6 and 7, we used an “A B B A A” design. From day 8 on, recordings were performed in the “A A B B A” and the “A B B A A” design in an alternating fashion. In all experiments, the object location sequence was kept as above. Recordings were preformed until the electrophysiological signal indicated that layer 1 was passed and no more cells could be recorded.

To obtain an unbiased estimation of the proportion of object cells in the targeted area of LEC, recordings began as soon as action potentials were detected for the first time. Tetrodes were gradually lowered by ~50 μm after each recording session with good coverage in all five trials, and only sessions with good coverage were included into the analysis. Recordings were continued until layer 1 was reached and the location of tetrode locations was histologically confirmed after the experiment. During the entire experiment, the experimenter was blind to the recorded cell types when advancing the tetrodes.

##### Two objects in context experiment (Figure 8)

Mice with tetrodes implanted into the LEC (n = 3 mice) were trained in two 20-minute open field trials per day for 5 days in a black box (70 cm x 70 cm x 30 cm) which contained a Lego object (6 cm x 8 cm x 14 cm) located 20 cm away from the top-left corner. The number of open field trials was gradually increased from 2 to 5 trials in the next 10 days of training. Tetrodes were lowered until spikes could be isolated. As the aim of this experiment was to characterize object cells, recordings were preferentially performed when object cells could be detected, and only sessions with object cells were included into the analysis. Recordings consisted of five 20-minute open field trials separated by 10-minute rest box periods. Two different objects were utilized: an irregular prism-shaped Lego tower (6 cm x 8 cm x 14 cm) and a coffee mug (8 cm in diameter, 10 cm in height). Each object was placed at a distance of 20 cm from one of the open field corners. The arrangement of objects is illustrated in Figure 8C. Two different spatial configurations of objects were examined (see Figure 8C).

#### Histology

After the electrophysiological recordings, mice were deeply anesthetized with ketamine and xylazine, and transcardially perfused first with phosphate buffered saline (PBS) and then fixed with 4% formaldehyde (Histofix, Roti). The brain was removed and maintained in 4% formaldehyde for at least 24h at 4ºC. Brains were then sliced coronally in 50 μm sections using a vibratome, mounted in Superfrost Plus™ slides (Thermo scientific) and stained with Cresyl violet to determine the location of the tetrode tracks. For the mice with a virus injection and tetrodes in the MS, alternating serial sections were stained with Cresyl violet and DAPI. Correct viral expression was assessed by examining the virus mediated mCherry expression in the DAPI stained sections. Imaging of the sections was performed using a slide scanner (Zeiss Axio Scan Z1).

#### Data analysis

All analysis was performed using custom R and MATLAB scripts. Basic electrophysiological properties such as spike waveform properties, mean firing rates and spatial properties were calculated using the ‘relectro’ R package (https://github.com/kevin-allen/relectro).

##### Spike sorting

The recorded electrophysiological signal was band-pass filtered (0.8-5 kHz) offline and spikes from individual neurons were subsequently extracted from this signal. Spikes were assigned to different clusters (i.e., putative neurons) via automatic clustering software (https://github.com/klusta-team/klustakwik) and then refined manually using the “Klusters” graphical interface software (http://neurosuite.sourceforge.net/). Clusters were only kept if spikes were detected in all open field trials and/or if spikes were detected in all rest box trials. Analysis was performed only on active neurons, i.e., neurons with a mean firing rate above 0.1 Hz in all open field trials.

##### Spike waveform parameters

The mean spike waveform for each neuron was calculated from the bandpass filtered signal of the tetrode wire with the largest spike amplitude. The trough-to-peak latency and spike asymmetry of the mean waveform were calculated in a 3-ms time window centered on the minimum voltage value (i.e., the trough of the spike). Spike asymmetry was calculated as:

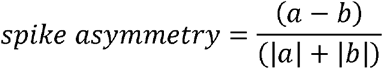

where a and b are the baseline-to-peak amplitudes of the mean spike waveform.

##### Post-processing of the tracking coordinates

All tracking data from open field trials were rescaled and aligned so that the bottom left corner of the open field box would have x-y coordinates (0,0), the top right corner of the boxes would have x-y coordinates (70,70) and the cue card and opening of the setup would be in the exact same position (left and bottom borders, respectively), before calculating the firing rate maps.

##### Firing rate maps

For each trial, the open field environment was divided in bins of two-by-two cm, and the mean firing rate of each neuron in that bin was calculated as the number of spikes divided by the time spent in that bin. The resulting map for each neuron was then smoothed using a Gaussian kernel function using a standard deviation of 3 cm. Only periods where mice were running at a speed ≥ 3 cm/s were used to calculate the firing rate maps.

##### Spatial information score

The spatial information score^68^ was calculated following the function:

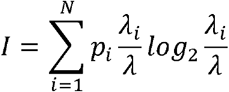

Where p_i_ is the occupancy probability of bin i in the firing rate map, λ_i_ is the firing rate of bin i and λ the mean firing rate of the neuron.

##### Field definition

Place field boundaries were determined from a joint reference map derived from the first and second open field trials and was performed as described previously^67,69^. Briefly, the firing rate map of each neuron was smoothed using a Gaussian filter and a place field was defined as a region in the smoothed firing rate map comprising ten or more contiguous bins where the bin’s firing rate exceeded 10% of the peak bin rate. Neurons with peak bin rates lower than 1.5 Hz were not assigned place fields. Field boundaries were drawn using MATLAB’s *contourc()* function. If any remaining bins outside of the place field surpassed the peak bin rate threshold, the procedure was repeated, and this was considered a new place field. The number of place fields of each neuron was defined as the number of independent, non-overlapping contours obtained with the *contourc()* function in one neuron’s firing rate map.

##### Map stability

The maps stability of each neuron was calculated as the correlation coefficient of the twodimensional matrices of two firing rate maps. The within-trial map stability was calculated between the first and second half of each open field trial (to determine whether a neuron was spatial) and the across-trial map stability was calculated between consecutive open field trials (to determine whether there was remapping or not).

##### Map stability after rotation of the rate maps

To evaluate whether the observed remapping was a result of the rotation of the firing rate map, one firing rate map was rotated analytically in steps of 10º (using the ‘firingRateMapsRotation’ function from the R package ‘relectro’) and the correlation coefficient between the rotated map and the non-rotated map was calculated for each step. In total the map was rotated by 10-350°, in 10° steps.

##### z-score

To calculate how selective a neuron’s firing was to the position of an object, we calculated the z-score analogous to what was described in ref.^28^, by following the formula:

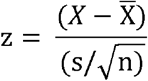

where X is the mean firing rate of the neuron in the 22 × 22 cm area that contains and surrounds the object, 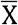 is the mean rate in a matching number of randomly selected bins outside the object-surrounding area, s is the standard deviation of the mean rate outside of the object-surrounding area and n is the number of bins from the firing rate map that are in the object-surrounding area.

##### Firing rate change

The absolute firing rate change of a neuron between two consecutive open field trials was defined as:

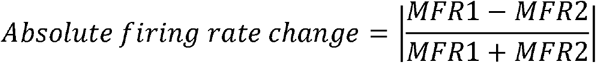

Where MFR1 and MFR2 are the mean firing rates of that neuron during open field trials 1 and 2, respectively.

##### Cell type classification

Isolated neurons were first divided into putative excitatory (mean firing rate in all open field trials > 0.1 and < 5.0 Hz, trough-to-peak latency > 0.4 ms and spike asymmetry < 0.1) or putative FS neurons (mean firing rate in all open field trials ≥ 5.0 Hz, trough-to-peak latency ≤ 0.4 ms and spike asymmetry ≥ 0.1). For the MEC recordings, we only included trough-locked theta-rhythmic neurons (obtained from ref.^67^ using the same selection as was described), which are typically found in the superficial layers of MEC^67^. Subsequently, neurons were deemed spatial or non-spatial. Spatial neurons had a spatial information score and within-trial map stability greater than the spatial information score and map stability obtained by chance in the first open field trial. For each neuron, chance levels were determined in a two-step process. First, we shuffled the position data 100 times and calculated the spatial information score and map stability for each iteration. This yielded a distribution of randomly generated values for each neuron. A neuron was considered spatial if its non-shuffled values exceeded the 95^th^ percentile of the randomly generated values for both the spatial information score and map stability. For the remapping experiments, neurons were included in the analysis if they were spatial in at least one context. For the object in context experiments, the same criteria were applied but, for each context, only trials with the object in the same position were considered. For the two objects in context experiment, neurons were included in the analysis if they were spatial in at least one of the five open field trials.

##### Classification of neurons into discriminating, stable and unstable (Figures 4 and 5)

Responses of spatially modulated neurons were divided in three groups based on the map stability and the discrimination index. The discrimination index was calculated as the difference between the median map stability between trials in the same box or room (W1 and W2) minus the median map stability between trials in different boxes or rooms (A1 and A2). Spatially modulated neurons were then categorized as discriminating (map stability in W1 and/or W2 > 0.4 and discrimination index > 0.25), stable (map stability in W1 and/or W2 > 0.4 and discrimination index < 0.25) or unstable (map stability in W1 and W2 < 0.4).

###### Classification of object and context coding neurons

In the object in context experiment (Figure 7), we used a z-score threshold of 2 around the object to categorize putative excitatory neurons in two types of object cells. Context-independent object cells were active around the object (z-score > 2) in the two positions within the same context and were also active around the object across different contexts (i.e., trials from sCdO and dCsO comparisons). Context-dependent object cells were active around the object in all trials in the black box, but not in the white box. Non-object cells were subdivided into spatially and non-spatially modulated neurons (see ‘Cell type classification’). Spatially modulated neurons were additionally subdivided in three groups: discriminating between contexts (i.e., context-coding neurons, sCsO map stability > 0.4 and dCsO map stability < 0.4), stable (sCsO map stability > 0.4 and dCsO map stability > 0.4) and unstable (sCsO map stability < 0.4). Finally, context-coding neurons were considered to remap to the object location if the their sCdO map stability was lower than 75% of their sCsO map stability.

###### Classification of object-coding neurons

For the recordings with two objects and two boxes (Figure 8), we used a z-score threshold of 2 around the objects to categorize putative excitatory neurons into the different categories of object cells. Context-dependent object identity cells were active around one specific object in all open field trials in one but not the other context (i.e., either in the first, second and fifth trial or in the third and fourth trial). Context-dependent object cells were active around both objects in all open field trials in one but not the other context (i.e., either in the first, second and fifth trial or in the third and fourth trial). Context-independent object-identity cells were active around the position of one of the two objects in both contexts (i.e., in the second and third trial). Context-independent object cells were active at the positions of both objects in both contexts (i.e., in the second and third trial).

##### Population vector analysis

For the remapping experiments, all discriminating neurons were used for the population vector analysis. The firing rate maps of all included neurons were stacked along the z-axis. The distribution of mean rates along the z-axis for a given x-y location (i.e., one bin of the firing rate map) represents the population vector for that bin. For each pair of trials, the Pearson’s correlation coefficient between the population vectors was calculated for each bin. Neurons with a peak bin rate < 1 Hz were excluded from this analysis. The correlation coefficients of all bins were averaged to estimate the median population vector correlation for a pair of sessions.

##### Place decoding from the population activity

Place decoding was performed using a maximum likelihood decoder of population spike count vectors as described in ref.^72^. In each open field trial from the recording sessions with more than 10 simultaneously recorded neurons, we computed place maps r_i_(x,y) for all neurons i = 1, … N separately. For decoding, we constructed population vectors of spike counts (k_1_, …, k_N_) in each T = 2 s interval and obtained the estimated (x,y) position by maximizing the Poisson loglikelihood ∑_i_ {k_i_ log[r_i_(x,y)T] – r_i_(x,y)T}. Finally, place decoding accuracy was assessed by calculating Pearson’s r between true and estimated coordinates.

### QUANTIFICATION AND STATISTICAL ANALYSIS

All statistical tests were performed with α = 0.05 and using the ‘rstatix’ package in R (v 0.7.0 https://CRAN.R-project.org/package=rstatix). All score distributions were tested for normality using the Shapiro-Wilk normality test. Because most of our datasets were not normally distributed, we performed non-parametric tests. All tests were two-sided. For comparison of scores across conditions, we performed Kruskal-Wallis ANOVA tests, if there were more than two comparisons (A1-A2-W1-W2 or sCdO-dCsO-sCsO-dCdO). Post-hoc comparisons were performed using Wilcoxon sum rank tests. Multiple comparisons were adjusted with a Holm-Bonferroni correction. For comparisons of the same neurons across two different trials we used Wilcoxon signed-rank tests. For comparisons between the two-box and the two-room experiments, we used Wilcoxon sum rank tests. To compare the proportions of neurons between groups, we used chi-square tests.

## Supplementary figures and figure legends

**Figure S1.**
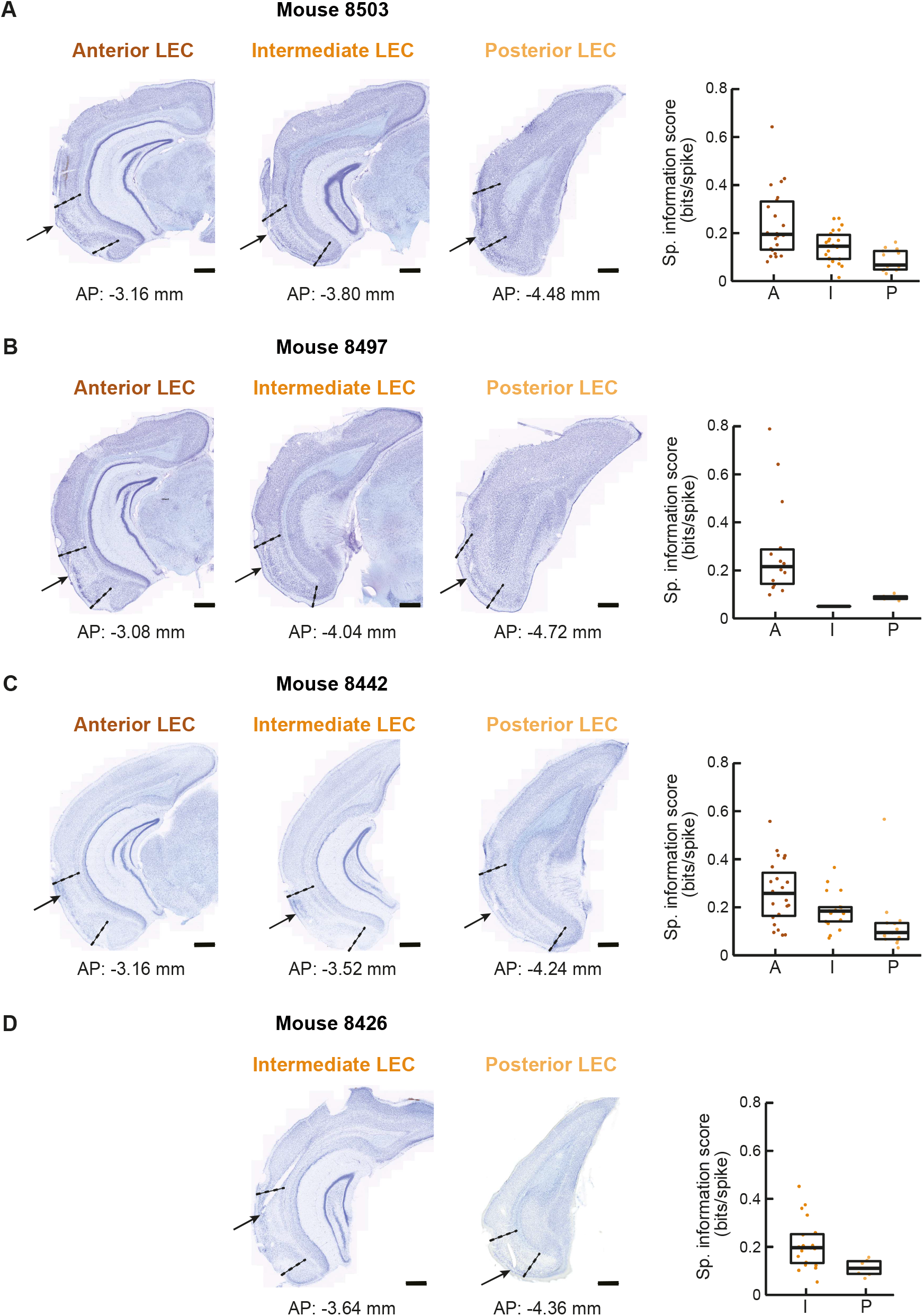
Tetrode locations in the LEC. Related to Figure 1. **(A-D)** Pictures of Cresyl violet-stained coronal sections of the brain of four mice subjected to recordings in the LEC to assess spatial selectivity of neurons along its antero-posterior axis. For each mouse, the histology pictures show the tetrode tracks at the anterior (left), intermediate (middle) and posterior LEC (right) positions. LEC borders are indicated with stippled lines and the tetrode tips are marked with black arrows. Antero-posterior coordinates in relation to bregma are specified underneath each picture. The tetrode in the anterior LEC coordinate from mouse ‘8426’ was broken, and was thus not included in the analysis. Boxplots depict medians and interquartile ranges of the spatial information scores of all excitatory neurons recorded in the tetrode at anterior (A), intermediate (I) and posterior (P) LEC coordinates for that mouse. Scale bar: 500 μm.

**Figure S2.**
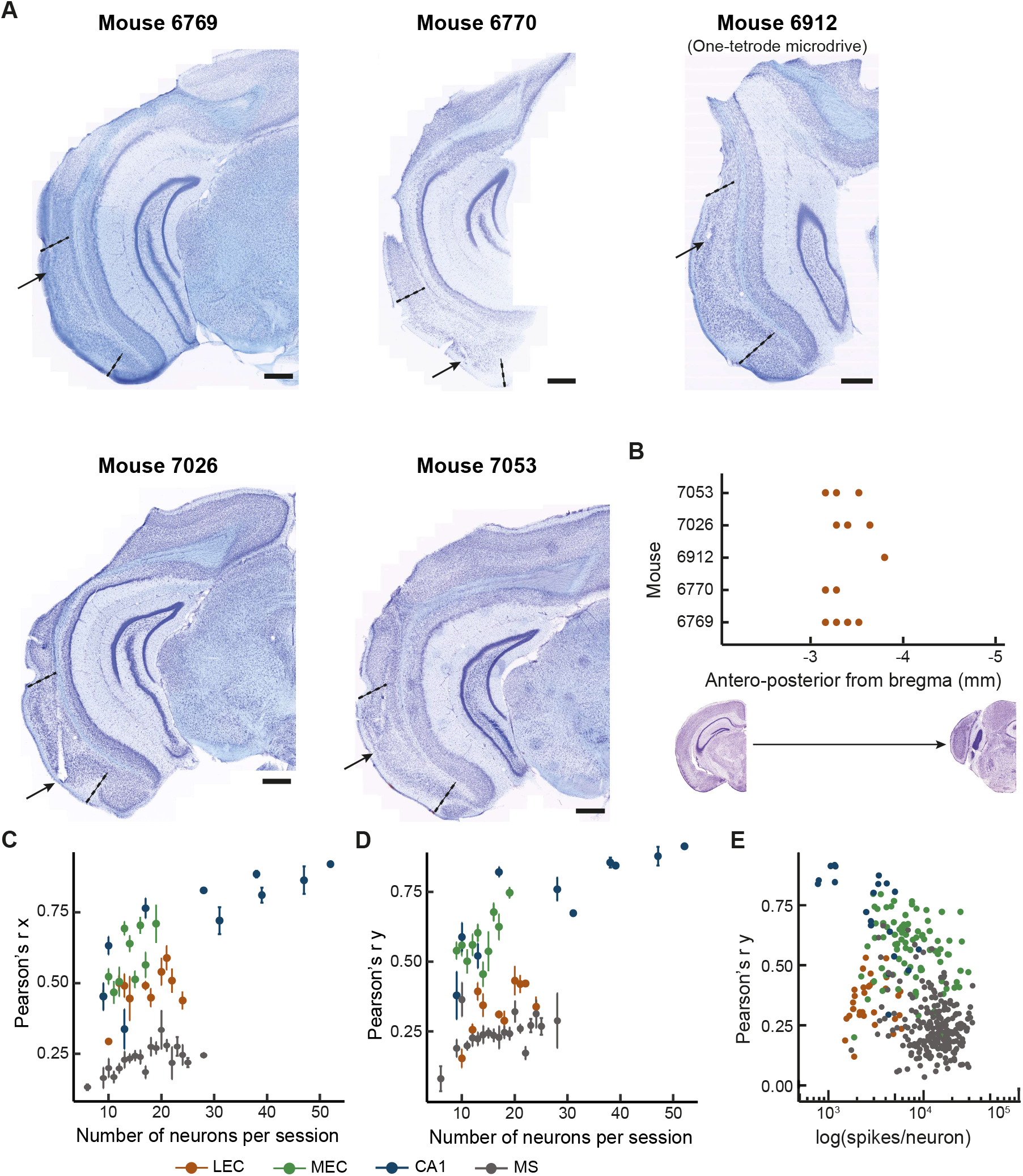
Tetrode locations in the anterior/intermediate LEC and additional place decoding analysis. Related to Figure 2. **(A)** Pictures of Cresyl violet-stained coronal sections from all LEC-implanted mice used for the analysis in Figure 2. For each mouse, one representative picture with a tetrode track in LEC is shown. LEC borders are outlined with stippled lines and the tetrode tips are indicated with black arrows. **(B)** Location of all visible tetrode tracks in the LEC antero-posterior axis in the five mice. Mouse ‘6912’ was implanted with a one-tetrode microdrive. **(C)** Pearson correlation (r) of the fitted x coordinate as a function of the number of neurons simultaneously recorded per session. Points show the median and error bars the standard deviation. **(D)** Same as **(C)**, but for the fitted y coordinate. **(E)** Pearson correlation (r) of the fitted y coordinate as a function of the mean spikes per neuron for each session. Scale bar: 500 μm

**Figure S3.**
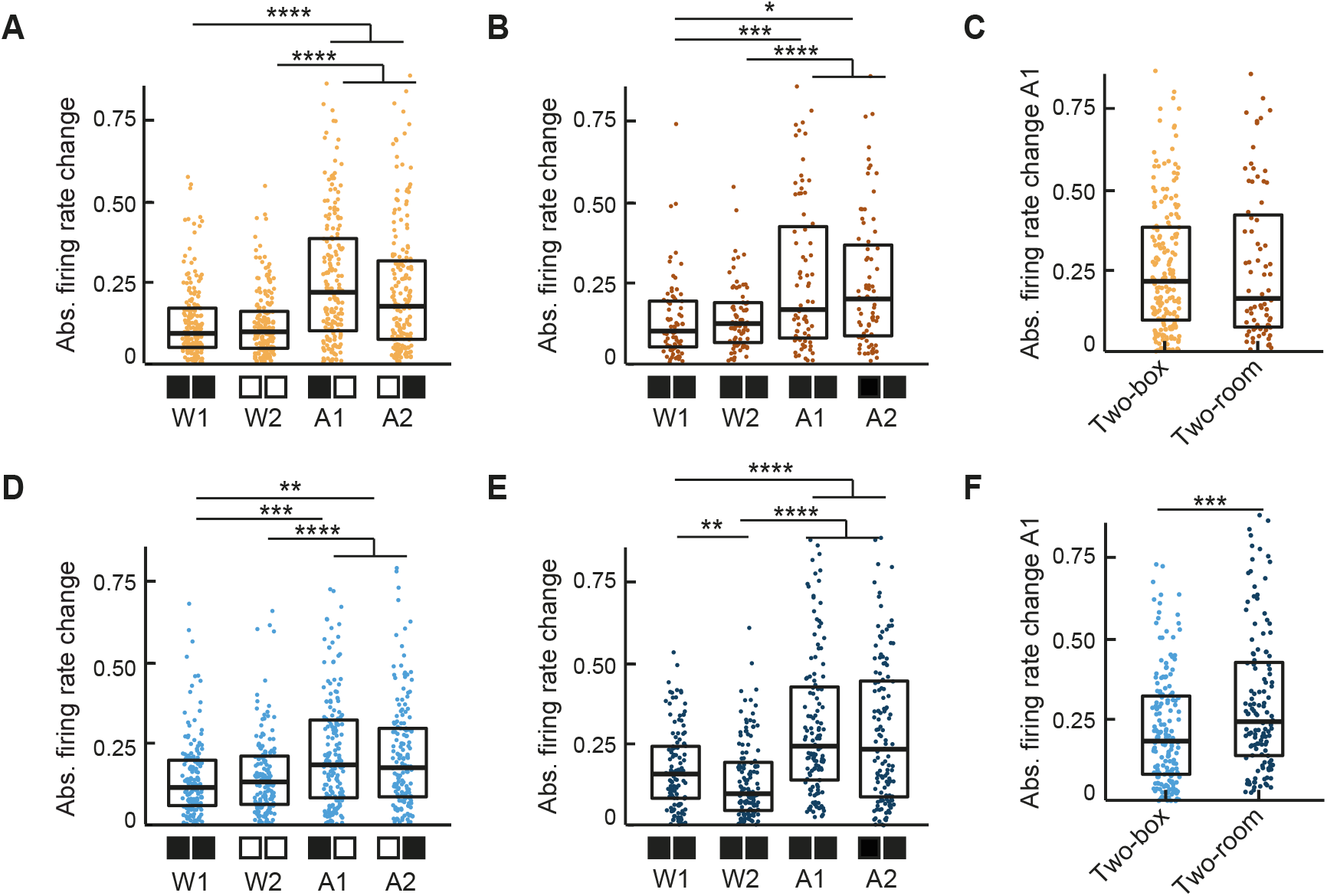
Firing rate change in LEC and CA1 neurons in the two-box and the two-room experiments. Related to Figures 4 and 5. **(A-B)** Mean firing rate change of discriminating excitatory spatially modulated neurons recorded in the LEC in the two-box **(A)** and the two-room **(B)** experiments. Absolute mean firing rate change is higher between trials in the two boxes and the two rooms (A1, A2), in the respective experiments, than between trials in the same box and in the same room (W1, W2) (both p < 0.0001, H = 74.794, df = 3, Kruskal-Wallis ANOVA, n neurons from m mice: two-box, 184 and 13; two-room, 92 and 5). **(C)** In LEC neurons, the mean firing rate change between trials in two different boxes is not significantly different from the change between trials in two different rooms (e.g., A1 in **(A)** vs. A1 in **(B))** (p > 0.05, W = 6731, Wilcoxon sum rank test). **(D-E)** Mean firing rate change of discriminating place cells recorded in CA1 in the two-box **(D)** and the two-room **(E)** experiments. Absolute mean firing rate change is higher between trials in the two boxes and the two rooms (A1, A2), in the respective experiments, than between trials in the same box and in the same room (W1, W2) (both p < 0.0001, H = 62.157, df = 3, Kruskal-Wallis ANOVA, n neurons from m mice: two-box, 167 and 4; two-room, 128 and 4). **(F)** In CA1 place cells, the mean firing rate change between trials in two different boxes is significantly different from the change between trials in two different rooms (p < 0.001, W = 13133, Wilcoxon signed-rank test). Boxplots show medians and interquartile ranges, and each point represents an individual neuron. Holm-Bonferroni correction procedure was applied for multiple comparisons. *p < 0.05, **p < 0.01, ***p < 0.001, ****p < 0.0001.

**Figure S4.**
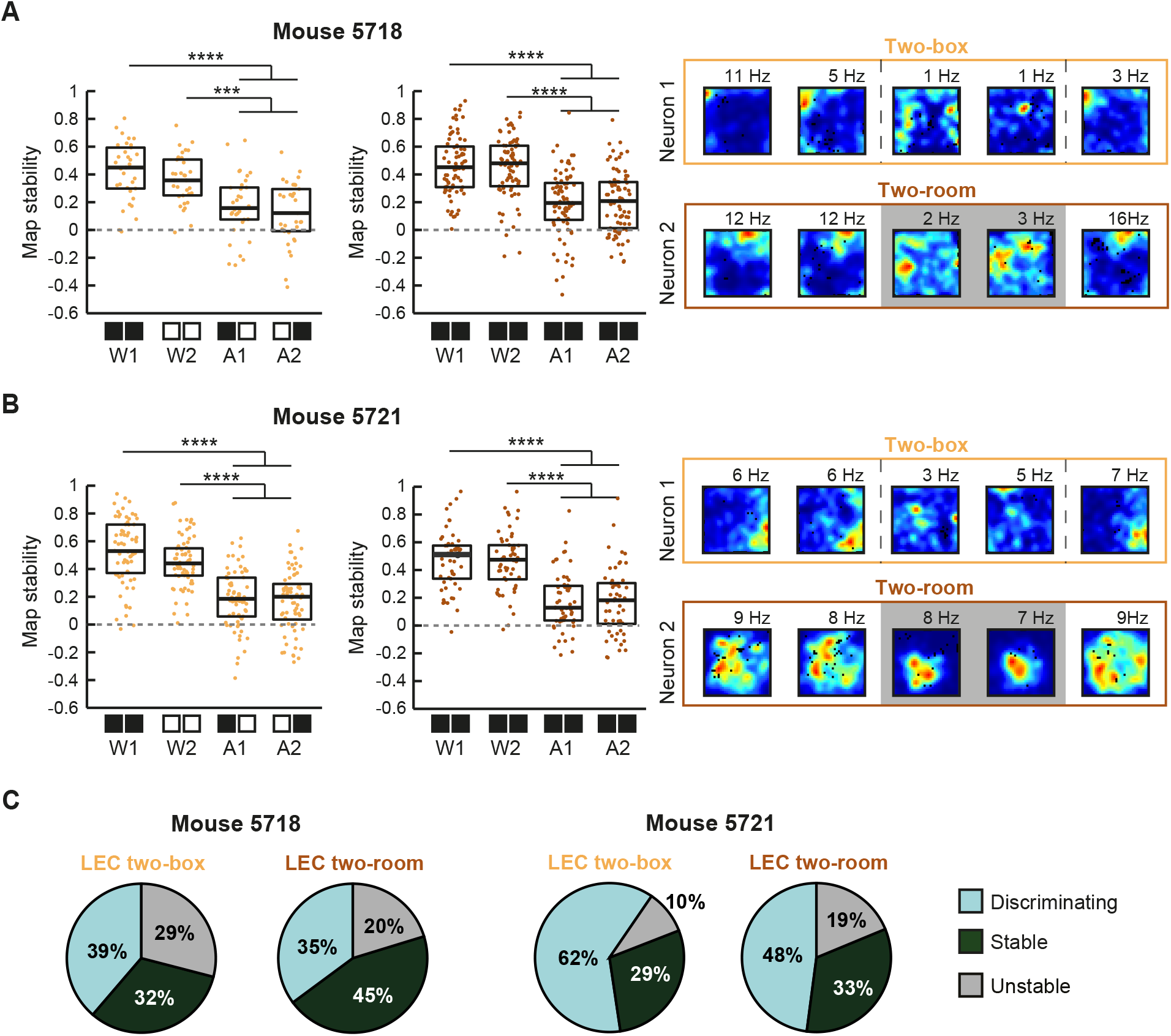
Remapping of LEC spatially modulated neurons in two mice subjected to both the two-box and the two-room experiments. Related to Figures 4 and 5. **(A-B)** Map stability of spatial LEC neurons recorded in the two-box experiment (yellow, left) and in the two-room experiment (middle, brown). Firing rate maps of an example neuron in each experimental setting are additionally shown (right). Panel **(A)** and **(B)** display data from mouse ‘5718’ (p < 0.05, H = 35.422, df = 3, Kruskal-Wallis ANOVA, n = 31 neurons in the two-box experiment; p < 0.0001, H = 77.056, df = 3, Kruskal-Wallis ANOVA, n = 74 neurons in the two-room experiment) and mouse ‘5721’ (p < 0.0001, H = 86.004, df = 3, Kruskal-Wallis ANOVA, n = 48 neurons in the two-box experiment; p < 0.001, H = 64.981, df = 3, Kruskal-Wallis ANOVA, n = 63 neurons in the two-room experiment), respectively. **(C)** Proportion of discriminating, stable and unstable neurons in the two-box (left) and the two-room (right) experimental settings for mouse ‘5718’. **(D)** Same as **(C)** for mouse ‘5721’. Boxplots represent median and interquartile range values, and each point the value for each individual neuron. Holm-Bonferroni correction was applied for multiple comparisons. ***p < 0.001, ****p < 0.0001.

**Figure S5.**
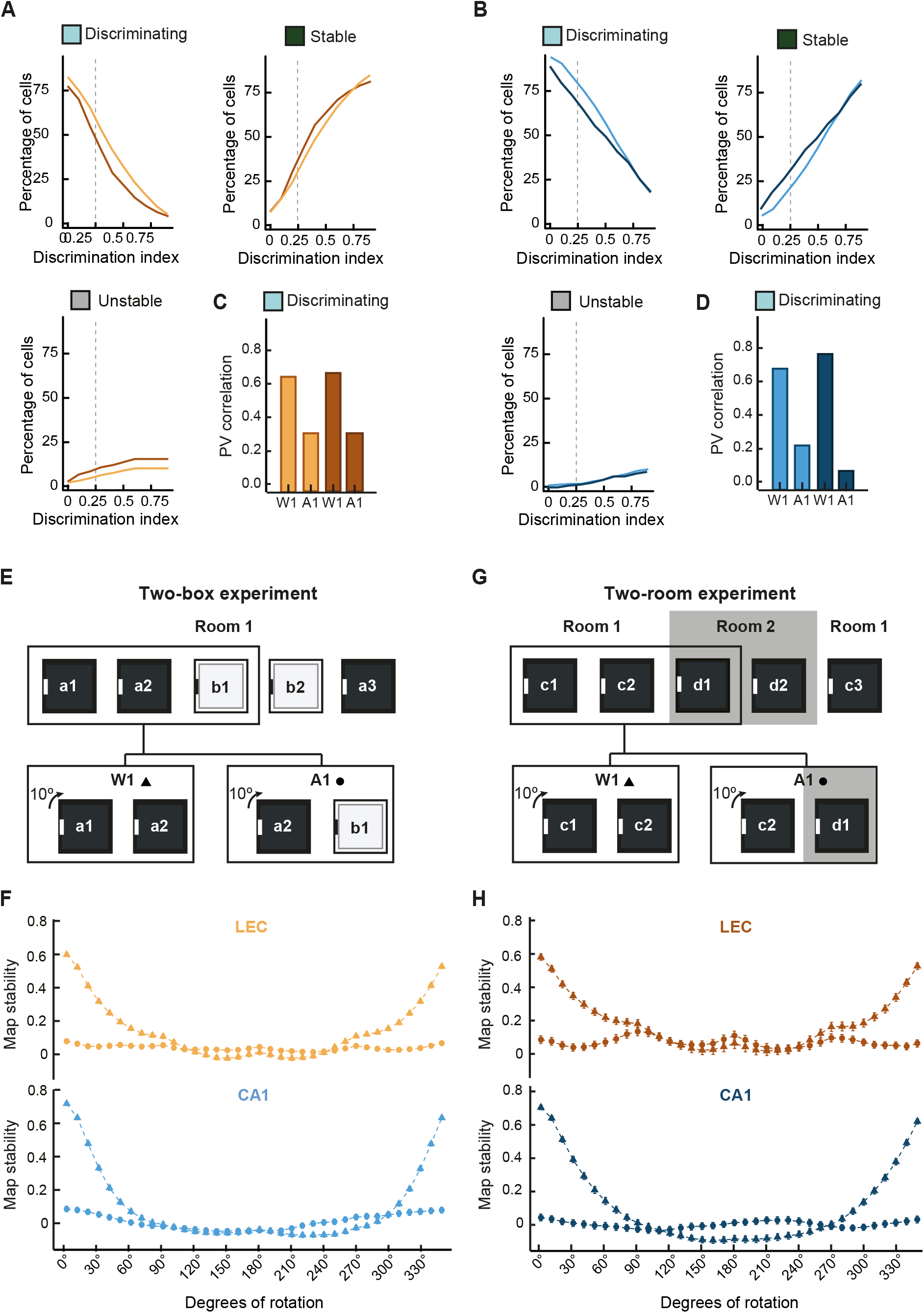
Remapping properties of LEC and CA1 spatially modulated neurons. Related to Figures 4 and 5. **(a)** Change of the proportion of discriminating, stable and unstable LEC spatially modulated neurons at different discrimination index thresholds in the two-box (yellow) and two-room (brown) experiments. **(B)** Same as in **(A)**, but for CA1 place cells recorded in the two-box (light blue) and two-room experimental setting (dark blue). **(C)** Population vector correlation of discriminating LEC neurons within and across contexts in the two-box (yellow) and the two-room (brown) experiments. **(D)** Same as **(C)**, but for CA1 place cells. **(E-G)** Remapping was not a result of a rotation of the map. **(E)** The firing rate maps from three consecutive open field trials (surrounded by a black square) were considered. For each neuron, map stability was calculated between the first and the second black box trial (a1 and a2) to assess within context map stability (W1), and between the second black box trial and the first white box trial (a2 and b1) to assess across context map stability (A1). One firing rate map was rotated in 10º steps and the map stability was calculated after each rotation. **(F)** Median map stability of LEC spatially modulated neurons (yellow, top) and CA1 place cells (light blue, bottom) at each rotation step in W1 (triangles) and A1 (circles) in the two-box experiment. **(G)** Same as **(E)** for the two-room experiment. For each neuron, map stability was calculated between two consecutive same-room trials (c1 and c2) to assess within context map stability (W1) and between the second trial in room 1 and the first trial in room 2 (c2 and d1) to assess across context map stability (A1). **(H)** Median map stability of LEC spatially modulated neurons (brown, top) and CA1 place cells (dark blue, bottom) at each rotation step in W1 (triangles) and A1 (circles) in the two-room experiment.

**Figure S6.**
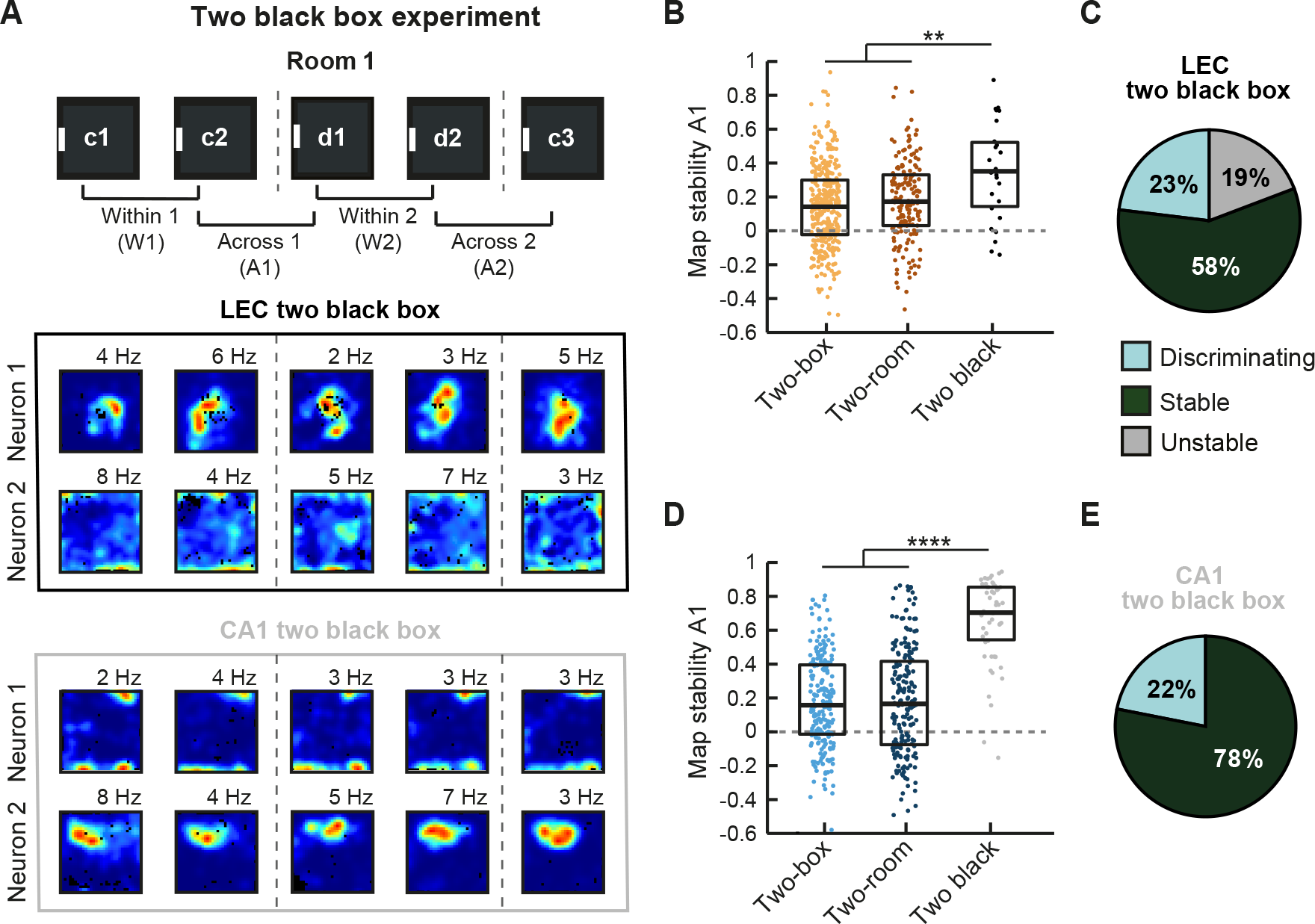
Absence of remapping in LEC and CA1 spatially modulated neurons when using two identical black boxes in the same location. Related to Figures 4 and 5. **(A)** (Top) Experimental setting: recordings were performed by alternating between two identical black boxes (c and d) placed in the same position in the same room. (Bottom) Color-coded firing rate maps from two example neurons recorded in LEC and in CA1 during the two-black box experimental setting (stippled lines denote the change between boxes). **(B)** Across boxes (A1), LEC spatially modulated neurons exhibit a higher map stability in the two-black box experiment compared to that in the two-box and the two-room experiments (p < 0.05, H = 9.6805, df = 2, Kruskal-Wallis ANOVA, n = 22 neurons from 4 mice in the two-black box experiment). **(C)** Proportion of discriminating, stable and unstable LEC neurons in the two-black box experiment. Note that the majority of LEC neurons remain stable across the two black boxes. **(D)** Map stability is also higher in the two-black box experiment in CA1 place cells (p < 0.0001, H = 88.929, df = 2, Kruskal-Wallis ANOVA, n = 52 neurons in the two-black box experiment). **(E)** Same as **(C)**, but for CA1 place cells. Boxplots represent median and interquartile range values, and each point the value for each individual neuron. Holm-Bonferroni correction was applied for multiple comparisons. **p < 0.01, ****p < 0.0001.

**Figure S7.**
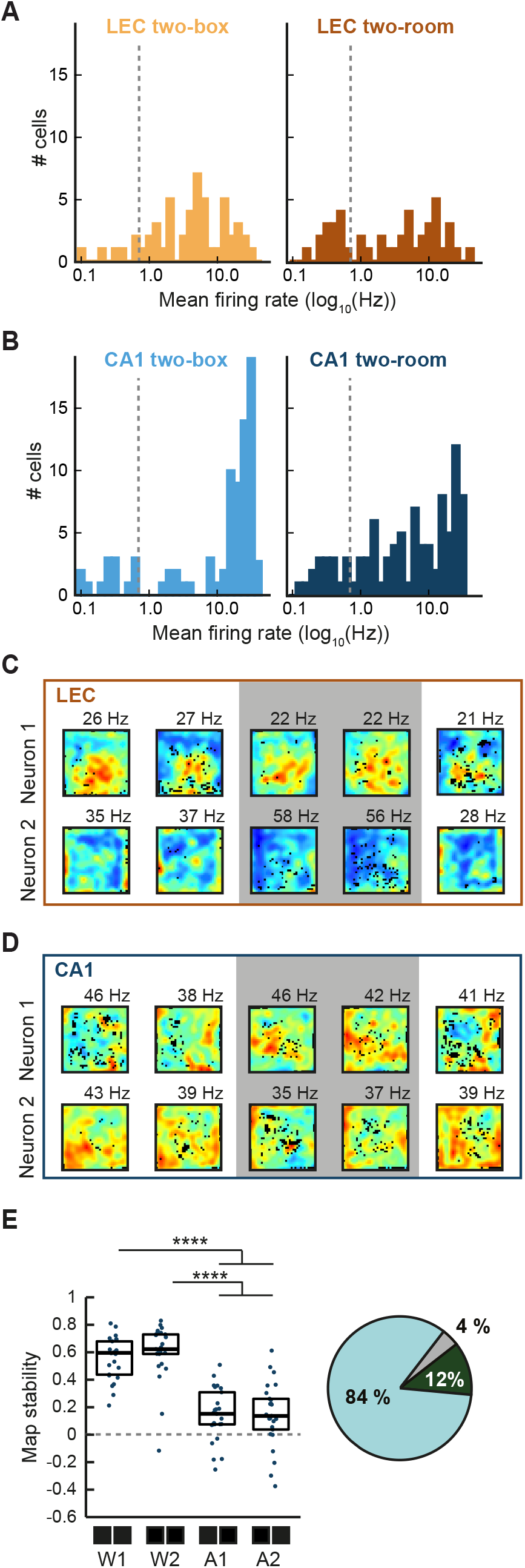
Selection and remapping of FS neurons in the two-room experiment. Related to Figure 6. **(A)** Distribution of mean firing rates of LEC neurons in the two-box (left) and the two-room (right) experiments that meet the spike waveform criteria of FS neurons as defined in Figure 3A. In addition, the cut-off for the mean firing rate was 5 Hz (stippled line) in all open field trials. **(B)** Same as in **(A)** for CA1 neurons recorded in the two-box (left) and two-room (right) experiments. **(C)** Color-coded rate maps of discriminating LEC FS neurons recorded in the two-room experiment. Grey background indicates the room change. **(D)** Same as **(C)** for two CA1 FS neurons. **(E)** (Left) The map stability of CA1 FS spatially modulated neurons is higher between trials in the same room (W1, W2) than between trials in different rooms (A1, A2) (p < 0.0001, H = 57.542, df = 3, Kruskal-Wallis ANOVA, n = 25 neurons from 4 mice). (Right) Proportion of discriminating, stable or unstable FS neurons using the same thresholds as for putative excitatory neurons. Note that in CA1 the vast majority of FS neurons discriminate between the two rooms. Boxplots represent median and interquartile range values, and each point the value for each individual neuron. Holm-Bonferroni correction procedure was applied for multiple comparisons. ****p ≤ 0.0001.

