## Supplementary material for "Distinct spatial maps and multiple object codes in the lateral entorhinal cortex": Key resources table

| **REAGENT or RESOURCE** | **SOURCE** | **IDENTIFIER** |
| --- | --- | --- |
| **Bacterial and virus strains** | | |
| pAAV-EF1a-double floxed-hChR2(H134R)-  mCherry-WPRE-HGHpA | Addgene | Cat# 20297 |
| **Chemicals, peptides, and recombinant proteins** | | |
| Cresyl violet acetate | Sigma | CAS: 10510-54-0 |
| Histofix 4% formaldehyde | Roti | Cat# 5666.2 |
| 4’,6-diamidino-2-phenylindole (DAPI) | Thermoscientific | Cat# 62248 |
| Mowiol® 40-88 | Sigma Aldrich | CAS: 9002-89-5 |
| EUKITT® mounting medium | Sigma Aldrich | CAS: 25608-33-7 |
| **Experimental models: Organisms/strains** | | |
| Mouse: wild type (C57BL/6N background) | N/A | N/A |
| Mouse: PV^Cre^ (C57BL/6N background) | (Fuchs et al., 2007) | N/A |
| **Deposited data** |  |  |
| CA1 dataset from Figures 2 and 3 | (Gil et al., 2018) | N/A |
| MEC dataset from Figures 2 and 3 | (Schlesiger et al., 2021) | N/A |
| **Software and algorithms** | | |
| R v 4.0.3 | <https://www.R-project.org/> | N/A |
| MATLAB v 2022a | Mathworks | RRID: SCR_001622 |
| Chronux v 2.12 v03 | <http://chronux.org/> | RRID: SCR_005547 |
| Relectro package v 0.0.0.9002 | <https://github.com/kevin-allen/relectro> | N/A |
| ‘rstatix’ package v 0.7.0 | <https://CRAN.R-project.org/package=rstatix> | RRID: SCR_021240 |
| ‘positrack’ position software | <https://github.com/kevin-allen/positrack> | N/A |
| ‘ktan’ acquisition software for the Intan Evaluation Board | <https://github.com/kevin-allen/ktan> | N/A |
| KlustaKwik | <https://github.com/klusta-team/klustakwik>;  (Kenneth et al., 2000) | RRID:SCR_014480 |
| Klusters | (Hazan et al., 2006) | RRID:SCR_008020 |
| Laser_stimulation | http://github.com/kevin-allen/laser_stimulation | N/A |
| **Other** | | |
| 4-tetrode microdrive | Axona | MDR-16TSS1 |
| 12-tetrode microdrive | Axona | MDR-48KDS1 |
| Tungsten tetrode wire 0.0005 mm | California Fine Wire Company | EW-12T |
| SST-33 tetrode tubes 80mm | Bilaney Consultants | SST-33/spc |
| Intan RHD2000 evaluation board | Intan Technologies | RHD2000 |
| 16-Channel Amplifier Board | Intan Technologies | RHD2216 |
| 64-Channel Amplifier Board | Intan Technologies | RHD2164 |
| Ain-76A Rodent Tablet 5 mg | TestDiet | 1813712 (5TUL) |
