## Supplementary material for "Distinct spatial maps and multiple object codes in the lateral entorhinal cortex": Statistics and quantification tables

**Supplementary information**

**Supplementary Table 1. Means, medians and standard error of the mean of all values reported in the manuscript.**

| **Parameter** | **Group** | **Mean** | **SEM** | **Median** |
| --- | --- | --- | --- | --- |
| Information score of excitatory cells along the antero-posterior axis | | | | |
|  | LEC_A | 0.2681258 | 0.02101406 | 0.2083967 |
|  | LEC_I | 0.1782892 | 0.01259848 | 0.1638741 |
|  | LEC_P | 0.1143751 | 0.0071466 | 0.1002868 |
| Number of fields along the AP axis | | |  |  |
|  | LEC_A | 3.791667 | 0.5433009 | 2 |
|  | LEC_I | 5.162791 | 0.6313429 | 4 |
|  | LEC_P | 5.866667 | 0.5150101 | 6 |
| Information score of excitatory cells in different areas | | | | |
|  | LEC | 0.3238187 | 0.01683741 | 0.2439353 |
|  | MEC | 0.7155526 | 0.0276197 | 0.5811109 |
|  | CA1 | 1.125829 | 0.04631841 | 0.9591684 |
|  | MS | 0.1022494 | 0.01046331 | 0.05647594 |
| Map stability (within) of excitatory cells in different areas | | | | |
|  | LEC | 0.3760915 | 0.01713017 | 0.3639758 |
|  | MEC | 0.6678015 | 0.01210459 | 0.7003633 |
|  | CA1 | 0.6207699 | 0.01639186 | 0.6711186 |
|  | MS | 0.1677504 | 0.01030158 | 0.1389478 |
| Pearson's r for estimated vs. true x position | | | |  |
|  | LEC | 0.4812085 | 0.01672282 | 0.472277 |
|  | MEC | 0.5360435 | 0.01451744 | 0.5414646 |
|  | CA1 | 0.7209447 | 0.04287851 | 0.791502 |
|  | MS | 0.2336738 | 0.00718008 | 0.2167513 |
| Pearson's r for estimated vs. true y position | | | |  |
|  | LEC | 0.3482081 | 0.01702998 | 0.3582313 |
|  | MEC | 0.5598193 | 0.01453523 | 0.5855796 |
|  | CA1 | 0.723587 | 0.03965891 | 0.8024636 |
|  | MS | 0.2449034 | 0.00743825 | 0.2269378 |
| Root-mean square estimation error | | | |  |
|  | LEC | 34.66688 | 0.4411188 | 35.15016 |
|  | MEC | 28.61945 | 0.4554686 | 28.27567 |
|  | CA1 | 22.72011 | 1.7034120 | 20.14325 |
|  | MS | 38.4239 | 0.2052631 | 38.77951 |
| Spikes per number of Neurons | | |  |  |
|  | LEC | 3.156.812 | 250.81 | 2.594.648 |
|  | MEC | 9377.456 | 668.263 | 7508.231 |
|  | CA1 | 3388.645 | 481.6637 | 3208.092 |
|  | MS | 14599.700 | 437.139 | 13992.340 |
| Mean firing rates of FS neurons in different areas | | | | |
|  | LEC | 20.880930 | 2.473459 | 20.844000 |
|  | MEC | 25.305730 | 0.994088 | 21.745940 |
|  | CA1 | 27.309990 | 4.250105 | 22.834870 |
|  | MS | 24.003950 | 0.893577 | 18.927530 |
| Map stability (across) of FS neurons in different areas | | | | |
|  | LEC | 0.5401472 | 0.0661168 | 0.517932 |
|  | MEC | 0.4962336 | 0.01346827 | 0.5379532 |
|  | CA1 | 0.7018117 | 0.03693461 | 0.716781 |
|  | MS | 0.2814533 | 0.00857201 | 0.2792935 |
| 2-box experiment LEC: map stability (across) of spatial excitatory neurons | | | | |
|  | W1 | 0.5184318 | 0.01281127 | 0.5276649 |
|  | W2 | 0.4609131 | 0.01337111 | 0.4621834 |
|  | A1 | 0.147371 | 0.01369749 | 0.1466754 |
|  | A2 | 0.1690802 | 0.0138423 | 0.1689234 |
| 2-room experiment LEC: map stability (across) of spatial excitatory neurons | | | | |
|  | W1 | 0.4876925 | 0.01684954 | 0.5056121 |
|  | W2 | 0.4792557 | 0.01835306 | 0.4897102 |
|  | A1 | 0.1823842 | 0.01861394 | 0.1767814 |
|  | A2 | 0.1853405 | 0.01876416 | 0.175646 |
| 2-box experiment CA1: map stability (across) of spatial excitatory neurons | | | | |
|  | W1 | 0.712615 | 0.01308935 | 0.7615878 |
|  | W2 | 0.6649484 | 0.01516642 | 0.7248006 |
|  | A1 | 0.1706746 | 0.01873687 | 0.1523691 |
|  | A2 | 0.1977482 | 0.01949029 | 0.1475731 |
| 2-room experiment CA1: map stability (across) of spatial excitatory neurons | | | | |
|  | W1 | 0.656835 | 0.01770309 | 0.7296187 |
|  | W2 | 0.6494406 | 0.01717502 | 0.7146773 |
|  | A1 | 0.1614987 | 0.02324621 | 0.148182 |
|  | A2 | 0.154298 | 0.02375172 | 0.1115383 |
| 2-box experiment LEC: absolute mean firing rate change of discriminating neurons | | | | |
|  | W1 | 0.1404869 | 0.00760582 | 0.1040287 |
|  | W2 | 0.130143 | 0.00632754 | 0.1014183 |
|  | A1 | 0.2593303 | 0.01214866 | 0.2125411 |
|  | A2 | 0.229903 | 0.01171191 | 0.1730606 |
| 2-room experiment LEC: absolute mean firing rate change of discriminating neurons | | | | |
|  | W1 | 0.164393 | 0.01148302 | 0.1221304 |
|  | W2 | 0.143285 | 0.00934257 | 0.1199216 |
|  | A1 | 0.2524861 | 0.01779213 | 0.170031 |
|  | A2 | 0.235135 | 0.0162793 | 0.180174 |
| 2-box experiment CA1: absolute mean firing rate change of discriminating neurons | | | | |
|  | W1 | 0.1526384 | 0.00878906 | 0.1235228 |
|  | W2 | 0.1417568 | 0.00801428 | 0.1193447 |
|  | A1 | 0.2147251 | 0.01170712 | 0.1815413 |
|  | A2 | 0.1996696 | 0.01120068 | 0.1629454 |
| 2-room experiment CA1: absolute mean firing rate change of discriminating neurons | | | | |
|  | W1 | 0.1730498 | 0.00912413 | 0.1517865 |
|  | W2 | 0.1358966 | 0.00853329 | 0.09746474 |
|  | A1 | 0.287048 | 0.01611438 | 0.2210339 |
|  | A2 | 0.2668309 | 0.01585514 | 0.2085718 |
| 2-box experiment LEC: PV correlation of discriminating neurons | | | | |
|  | W1 | 0.6848 | 0.0026 | 0.6893 |
|  | W2 | 0.7408 | 0.0027 | 0.7386 |
|  | A1 | 0.3025 | 0.0042 | 0.296 |
|  | A2 | 0.3425 | 0.0047 | 0.3379 |
| 2-room experiment LEC: PV correlation of discriminating neurons | | | | |
|  | W1 | 0.6698 | 0.0028 | 0.6752 |
|  | W2 | 0.7663 | 0.002 | 0.764 |
|  | A1 | 0.3026 | 0.0039 | 0.3057 |
|  | A2 | 0.3119 | 0.0039 | 0.3162 |
| 2-box experiment CA1: PV correlation of discriminating neurons | | | | |
|  | W1 | 0.7821 | 0.027 | 0.7935 |
|  | W2 | 0.7965 | 0.0026 | 0.8071 |
|  | A1 | 0.264 | 0.0037 | 0.2529 |
|  | A2 | 0.2936 | 0.0032 | 0.2887 |
| 2-room experiment CA1: PV correlation of discriminating neurons | | | | |
|  | W1 | 0.7303 | 0.0032 | 0.7424 |
|  | W2 | 0.797 | 0.0022 | 0.8087 |
|  | A1 | 0.1372 | 0.0028 | 0.1247 |
|  | A2 | 0.0958 | 0.0027 | 0.0802 |
| 2-black-box experiment LEC: map stability (across) of spatial excitatory neurons | | | | |
|  | W1 | 0.4722983 | 0.05583463 | 0.4639429 |
|  | W2 | 0.48542 | 0.05007528 | 0.4967406 |
|  | A1 | 0.3121331 | 0.05706134 | 0.3363646 |
|  | A2 | 0.3211824 | 0.04793658 | 0.2689542 |
| 2-black-box experiment CA1: map stability (across) of spatial excitatory neurons | | | | |
|  | W1 | 0.7644565 | 0.0255467 | 0.8136154 |
|  | W2 | 0.773832 | 0.0157401 | 0.7930162 |
|  | A1 | 0.6659136 | 0.03213793 | 0.7001648 |
|  | A2 | 0.7135298 | 0.02718407 | 0.7757793 |
| 2-box experiment LEC: map stability (across) of spatial FS neurons | | | | |
|  | W1 | 0.6096857 | 0.03837886 | 0.6241098 |
|  | W2 | 0.5643399 | 0.03952134 | 0.5410357 |
|  | A1 | 0.3002207 | 0.08036924 | 0.2743964 |
|  | A2 | 0.31391 | 0.0732578 | 0.322548 |
| 2-room experiment LEC: map stability (across) of spatial FS neurons | | | | |
|  | W1 | 0.6250155 | 0.06514474 | 0.6236273 |
|  | W2 | 0.5933881 | 0.06387906 | 0.5963743 |
|  | A1 | 0.4432641 | 0.07359521 | 0.4673683 |
|  | A2 | 0.3812862 | 0.0641259 | 0.3717874 |
| 2-box experiment CA1: map stability (across) of spatial FS neurons | | | | |
|  | W1 | 0.5886635 | 0.01902831 | 0.5978872 |
|  | W2 | 0.5978317 | 0.02281028 | 0.5885117 |
|  | A1 | 0.1485658 | 0.02655706 | 0.1436366 |
|  | A2 | 0.1062212 | 0.02601183 | 0.1186224 |
| 2-room experiment CA1: map stability (across) of spatial FS neurons | | | | |
|  | W1 | 0.558982 | 0.03085925 | 0.5977035 |
|  | W2 | 0.6019402 | 0.04194678 | 0.6247831 |
|  | A1 | 0.1418713 | 0.03894305 | 0.1492566 |
|  | A2 | 0.1412797 | 0.04744714 | 0.1338007 |
| 1object-2box experiment LEC: map stability (across) of discriminating context-coding neurons | | | | |
|  | sCsO | 0.4557211 | 0.0278046 | 0.465822 |
|  | sCdO | 0.4730822 | 0.02926075 | 0.5036489 |
|  | dCsO | 0.0861751 | 0.02089039 | 0.08558765 |
|  | dCdO | 0.09685055 | 0.02207246 | 0.06724213 |
| 1object-2box experiment CA1: map stability (across) of discriminating context-coding neurons | | | | |
|  | sCsO | 0.7034017 | 0.02170423 | 0.7668602 |
|  | sCdO | 0.4861052 | 0.0284675 | 0.5186243 |
|  | dCsO | 0.09808834 | 0.01923178 | 0.09622735 |
|  | dCdO | 0.1134971 | 0.02418896 | 0.0858727 |
| sCdO-sCdO difference of discriminating context-coding neurons | | | | |
|  | LEC | -0.0173611 | 0.04468596 | 0.02330323 |
|  | CA1 | 0.2172965 | 0.03397066 | 0.2170119 |

**Supplementary Table 2. Statistics from all tests performed in the main figures.**

| **Comparison** | **Statistical test** | **Test value** | **p-value** | **neurons (n)** | **Mice (N)** |
| --- | --- | --- | --- | --- | --- |
| Information score a1 AP gradient | Kruskal-Wallis test | H = 63.726; df = 2 | 1.45E-14 | n = 113 anterior LEC; n = 92 intermediate LEC; n = 90 posterior LEC | 5 |
| LEC A vs. LEC I | Wilcoxon sum rank test with Bonferroni correction | 6777 | 5.61E-04 |  | 4 vs. 5 |
| LEC A vs. LEC P | Wilcoxon sum rank test with Bonferroni correction | 8332 | 1.75E-14 |  | 4 vs. 5 |
| LEC I vs. LEC P | Wilcoxon sum rank test with Bonferroni correction | 5708 | 3.09E-05 |  | 5 vs. 5 |
| Number of fields AP gradient | Kruskal-Wallis test | H = 8.7103; df = 2 | 0.01284 | n = 113 anterior LEC; n = 92 intermediate LEC; n = 90 posterior LEC | 5 |
| LEC A vs. LEC I | Wilcoxon sum rank test with Bonferroni correction | 800 | 0.182 |  | 4 vs. 5 |
| LEC A vs. LEC P | Wilcoxon sum rank test with Bonferroni correction | 978 | 0.011 |  | 4 vs. 5 |
| LEC I vs. LEC P | Wilcoxon sum rank test with Bonferroni correction | 1154 | 1 |  | 5 vs. 5 |
| Prop. excitatory spatial neurons (Figure 2) | 4-sample test for equality of proportions without continuity correction | X-squared = 531.66; df = 3 | < 2.2E-16 | LEC: 211; MEC: 298; CA1: 240; MS: 375 | LEC: 5; MEC: 10; CA1: 4; MS: 4 |
| LEC vs. CA1 | Pairwise comparison of proportions with Bonferroni correction | . | 5.20E-09 |  |  |
| LEC vs. MEC | Pairwise comparison of proportions with Bonferroni correction | . | < 2.2E-16 |  |  |
| LEC vs. MS | Pairwise comparison of proportions with Bonferroni correction | . | < 2.2E-16 |  |  |
| CA1 vs. MEC | Pairwise comparison of proportions with Bonferroni correction | . | 0.043 |  |  |
| CA1 vs. MS | Pairwise comparison of proportions with Bonferroni correction | . | < 2.2E-16 |  |  |
| MEC vs. MS | Pairwise comparison of proportions with Bonferroni correction | . | < 2.2E-16 |  |  |
| Information scores of excitatory neurons (Figure 2) | Kruskal-Wallis test | H = 748.84; df = 3 | < 2.2E-16 | LEC: 211; MEC: 298; CA1: 240; MS: 375 | LEC: 5; MEC: 10; CA1: 4; MS: 4 |
| LEC vs. CA1 | Wilcoxon sum rank test with Bonferroni correction | 5459 | 4.13E-46 |  |  |
| LEC vs. MEC | Wilcoxon sum rank test with Bonferroni correction | 12636 | 7.74E-30 |  |  |
| LEC vs. MS | Wilcoxon sum rank test with Bonferroni correction | 71516 | 1.54E-58 |  |  |
| CA1 vs. MEC | Wilcoxon sum rank test with Bonferroni correction | 22735 | 2.21E-12 |  |  |
| CA1 vs. MS | Wilcoxon sum rank test with Bonferroni correction | 88071 | 1.54E-88 |  |  |
| MEC vs. MS | Wilcoxon sum rank test with Bonferroni correction | 108275 | 2.33E-96 |  |  |
| Within-trial map stability of excitatory neurons in HPR areas | Kruskal-Wallis test | H = 540.35; df = 3 | < 2.2e-16 | LEC: 211; MEC: 298; CA1: 240; MS: 375 | LEC: 5; MEC: 10; CA1: 4; MS: 4 |
| LEC vs. CA1 | Wilcoxon sum rank test with Bonferroni correction | 12500 | 9.96E-20 |  |  |
| LEC vs. MEC | Wilcoxon sum rank test with Bonferroni correction | 11551 | 2.84E-33 |  |  |
| LEC vs. MS | Wilcoxon sum rank test with Bonferroni correction | 58928 | 4.40E-22 |  |  |
| CA1 vs. MEC | Wilcoxon sum rank test with Bonferroni correction | 38707 | 6.00E-01 |  |  |
| CA1 vs. MS | Wilcoxon sum rank test with Bonferroni correction | 81715 | 1.23E-64 |  |  |
| MEC vs. MS | Wilcoxon sum rank test with Bonferroni correction | 105153 | 2.38E-85 |  |  |
| Rx decoding | Kruskal-Wallis test | H = 218.06; df = 3 | < 2.2e-16 | LEC: 32; MEC: 88; CA1: 20; MS: 228 sessions | LEC: 4; MEC: 8; CA1: 4; MS: 4 |
| LEC vs. MEC | Wilcoxon sum rank test with Bonferroni correction | 997 | 0.109 |  |  |
| LEC vs. CA1 | Wilcoxon sum rank test with Bonferroni correction | 102 | 9.66E-05 |  |  |
| LEC vs. MS | Wilcoxon sum rank test with Bonferroni correction | 6880 | 8.70E-16 |  |  |
| MEC vs. CA1 | Wilcoxon sum rank test with Bonferroni correction | 356 | 2.44E-04 |  |  |
| MEC vs. MS | Wilcoxon sum rank test with Bonferroni correction | 18729 | 1.48E-34 |  |  |
| CA1 vs. MS | Wilcoxon sum rank test with Bonferroni correction | 4411 | 1.07E-11 |  |  |
| Ry decoding | Kruskal-Wallis test | H = 200.63; df = 3 | < 2.2e-16 | LEC: 32; MEC: 88; CA1: 20; MS: 228 sessions | LEC: 4; MEC: 8; CA1: 4; MS: 4 |
| LEC vs. MEC | Wilcoxon sum rank test with Bonferroni correction | 301 | 3.80E-10 |  |  |
| LEC vs. CA1 | Wilcoxon sum rank test with Bonferroni correction | 28 | 1.75E-09 |  |  |
| LEC vs. MS | Wilcoxon sum rank test with Bonferroni correction | 5648 | 1.63E-06 |  |  |
| MEC vs. CA1 | Wilcoxon sum rank test with Bonferroni correction | 387 | 6.90E-04 |  |  |
| MEC vs. MS | Wilcoxon sum rank test with Bonferroni correction | 18695 | 2.68E-34 |  |  |
| CA1 vs. MS | Wilcoxon sum rank test with Bonferroni correction | 4430 | 6.84E-12 |  |  |
| FS mean firing rates in a1 (Figure 3) | Kruskal-Wallis test | H = 6.6563; df = 3 | 0.0837 | LEC: 10; MEC: 220; CA1:13; MS: 401 | LEC: 5; MEC: 10; CA1: 4; MS: 4 |
| Proportion of FS spatial neurons (Figure 3) | 4-sample test for equality of proportions without continuity correction | X-squared = 153.49; df = 3 | < 2.2e-16 | LEC: 10; MEC: 220; CA1:13; MS: 401 | LEC: 5; MEC: 10; CA1: 4; MS: 4 |
| LEC vs. CA1 | Pairwise comparison of proportions with Bonferroni correction | . | 0.54 |  |  |
| LEC vs. MEC | Pairwise comparison of proportions with Bonferroni correction | . | 0.95 |  |  |
| LEC vs. MS | Pairwise comparison of proportions with Bonferroni correction | . | 5.6e-06 |  |  |
| CA1 vs. MEC | Pairwise comparison of proportions with Bonferroni correction | . | 0.39 |  |  |
| CA1 vs. MS | Pairwise comparison of proportions with Bonferroni correction | . | 0.32 |  |  |
| MEC vs. MS | Pairwise comparison of proportions with Bonferroni correction | . | < 2E-16 |  |  |
| 2-box LEC: map stability of spatial neurons | Kruskal-Wallis test | H = 425.55; df = 3 | < 2E-16 | 307 neurons | 13 mice |
| W1 vs. W2 | Wilcoxon signed-rank tests with Bonferroni correction | 28377 | 1.40E-02 |  |  |
| W1 vs. A1 | Wilcoxon signed-rank tests with Bonferroni correction | 45656 | 1.22E-44 |  |  |
| W1 vs. A2 | Wilcoxon signed-rank tests with Bonferroni correction | 45212 | 6.78E-43 |  |  |
| W2 vs. A1 | Wilcoxon signed-rank tests with Bonferroni correction | 43985 | 2.92E-38 |  |  |
| W2 vs. A2 | Wilcoxon signed-rank tests with Bonferroni correction | 44764 | 3.58E-41 |  |  |
| A1 vs. A2 | Wilcoxon signed-rank tests with Bonferroni correction | 20946 | 5.02E-01 |  |  |
| 2-room LEC: map stability of spatial neurons | Kruskal-Wallis test | H = 197.29; df = 3 | < 2E-16 | 160 neurons | 5 mice |
| W1 vs. W2 | Wilcoxon signed-rank tests with Bonferroni correction | 6417 | 1 |  |  |
| W1 vs. A1 | Wilcoxon signed-rank tests with Bonferroni correction | 12369 | 3.31E-23 |  |  |
| W1 vs. A2 | Wilcoxon signed-rank tests with Bonferroni correction | 12399 | 1.96E-23 |  |  |
| W2 vs. A1 | Wilcoxon signed-rank tests with Bonferroni correction | 12363 | 3.68E-23 |  |  |
| W2 vs. A2 | Wilcoxon signed-rank tests with Bonferroni correction | 12313 | 8.70E-23 |  |  |
| A1 vs. A2 | Wilcoxon signed-rank tests with Bonferroni correction | 6588 | 1 |  |  |
| Prop. Remapping types LEC 2-box vs. 2-room | 6-sample test for equality of proportions without continuity correction | X-squared = 207.27; df = 5 | < 2E-16 | 2-box: 307 ; 2-room: 160 | 2-box: 12; 2-room: 5 |
| Discriminating | 2-sample test for equality of proportions without continuity correction | X-squared = 6.7435; df = 1 | 0.009409 |  |  |
| Stable | 2-sample test for equality of proportions without continuity correction | X-squared = 2.2032; df = 1 | 0 |  |  |
| Unstable | 2-sample test for equality of proportions without continuity correction | X-squared = 2.769; df = 1 | 0 |  |  |
| 2-box CA1: map stability of spatial neurons | Kruskal-Wallis test | H = 446.09; df = 3 | < 2E-16 | 212 neurons | 4 mice |
| W1 vs. W2 | Wilcoxon signed-rank tests with Bonferroni correction | 13688 | 4.40E-02 |  |  |
| W1 vs. A1 | Wilcoxon signed-rank tests with Bonferroni correction | 22325 | 3.28E-34 |  |  |
| W1 vs. A2 | Wilcoxon signed-rank tests with Bonferroni correction | 22195 | 1.97E-33 |  |  |
| W2 vs. A1 | Wilcoxon signed-rank tests with Bonferroni correction | 22395 | 1.24E-34 |  |  |
| W2 vs. A2 | Wilcoxon signed-rank tests with Bonferroni correction | 21983 | 3.52E-32 |  |  |
| A1 vs. A2 | Wilcoxon signed-rank tests with Bonferroni correction | 9795 | 5.69E-01 |  |  |
| 2-room CA1: map stability of spatial neurons | Kruskal-Wallis test | H = 321.62; df = 3 | < 2E-16 | 181 neurons | 4 mice |
| W1 vs. W2 | Wilcoxon signed-rank tests with Bonferroni correction | 8685 | 1 |  |  |
| W1 vs. A1 | Wilcoxon signed-rank tests with Bonferroni correction | 15987 | 2.84E-27 |  |  |
| W1 vs. A2 | Wilcoxon signed-rank tests with Bonferroni correction | 15849 | 2.42E-26 |  |  |
| W2 vs. A1 | Wilcoxon signed-rank tests with Bonferroni correction | 15960 | 4.33E-27 |  |  |
| W2 vs. A2 | Wilcoxon signed-rank tests with Bonferroni correction | 15954 | 4.75E-27 |  |  |
| A1 vs. A2 | Wilcoxon signed-rank tests with Bonferroni correction | 8516 | 1 |  |  |
| Prop. Remapping types CA1 2-box vs. 2-room | 6-sample test for equality of proportions without continuity correction | X-squared = 505.36; df = 5 | < 2E-16 | 2-box: 212; 2-room: 181 | 2-box: 4; 2-room: 4 |
| Discriminating | 2-sample test for equality of proportions without continuity correction | X-squared = 2.9681; df = 1 | 0.08492 |  |  |
| Stable | 2-sample test for equality of proportions without continuity correction | X-squared = 2.1229; df = 1 | 0.1451 |  |  |
| Unstable | 2-sample test for equality of proportions without continuity correction | X-squared = 0.34157; df = 1 | 0.5589 |  |  |
| 2-box LEC: map stability of FS spatial neurons | Kruskal-Wallis test | H = 16.674; df = 3 | 0.0008245 | 12 neurons | 5 mice |
| W1 vs. W2 | Wilcoxon signed-rank tests with Bonferroni correction | 50 | 1 |  |  |
| W1 vs. A1 | Wilcoxon signed-rank tests with Bonferroni correction | 71 | 0.056 |  |  |
| W1 vs. A2 | Wilcoxon signed-rank tests with Bonferroni correction | 72 | 0.041 |  |  |
| W2 vs. A1 | Wilcoxon signed-rank tests with Bonferroni correction | 75 | 0.015 |  |  |
| W2 vs. A2 | Wilcoxon signed-rank tests with Bonferroni correction | 73 | 0.029 |  |  |
| A1 vs. A2 | Wilcoxon signed-rank tests with Bonferroni correction | 36 | 1 |  |  |
| 2-box CA1: map stability of FS spatial neurons | Kruskal-Wallis test | H = 125.99; df = 3 | < 2E-16 | 45 neurons | 4 mice |
| W1 vs. W2 | Wilcoxon signed-rank tests with Bonferroni correction | 477 | 1 |  |  |
| W1 vs. A1 | Wilcoxon signed-rank tests with Bonferroni correction | 1033 | 1.03E-12 |  |  |
| W1 vs. A2 | Wilcoxon signed-rank tests with Bonferroni correction | 1035 | 3.41E-13 |  |  |
| W2 vs. A1 | Wilcoxon signed-rank tests with Bonferroni correction | 1028 | 6.48E-12 |  |  |
| W2 vs. A2 | Wilcoxon signed-rank tests with Bonferroni correction | 1035 | 3.41E-13 |  |  |
| A1 vs. A2 | Wilcoxon signed-rank tests with Bonferroni correction | 649 | 0.84 |  |  |
| 2box-1object LEC: map stability of discriminating context-coding neurons | Kruskal-Wallis test | H = 122.06; df = 3 | < 2E-16 | 134 neurons in total / 64 neurons discriminating | 6 mice |
| sCsO vs. sCdO | Wilcoxon signed-rank tests with Bonferroni correction | 998 | 1 |  |  |
| sCsO vs. dCsO | Wilcoxon signed-rank tests with Bonferroni correction | 2047 | 1.01E-10 |  |  |
| sCsO vs. dCdO | Wilcoxon signed-rank tests with Bonferroni correction | 2023 | 3.01E-10 |  |  |
| sCdO vs. dCsO | Wilcoxon signed-rank tests with Bonferroni correction | 2006 | 6.42E-10 |  |  |
| sCdO vs. dCdO | Wilcoxon signed-rank tests with Bonferroni correction | 1961 | 4.48E-09 |  |  |
| dCsO vs. dCdO | Wilcoxon signed-rank tests with Bonferroni correction | 1009 | 1 |  |  |
| 2box-1object CA1: map stability of discriminating context-coding neurons | Kruskal-Wallis test | H = 206.54; df = 3 | < 2E-16 | 175 neurons total// 95 neurons discriminating | 3 mice |
| sCsO vs. sCdO | Wilcoxon signed-rank tests with Bonferroni correction | 3746 | 3.20E-07 |  |  |
| sCsO vs. dCsO | Wilcoxon signed-rank tests with Bonferroni correction | 4554 | 1.92E-16 |  |  |
| sCsO vs. dCdO | Wilcoxon signed-rank tests with Bonferroni correction | 4509 | 7.92E-16 |  |  |
| sCdO vs. dCsO | Wilcoxon signed-rank tests with Bonferroni correction | 4275 | 7.98E-16 |  |  |
| sCdO vs. dCdO | Wilcoxon signed-rank tests with Bonferroni correction | 4230 | 2.77E-12 |  |  |
| dCsO vs. dCdO | Wilcoxon signed-rank tests with Bonferroni correction | 2199 | 1.00E+00 |  |  |
| 2box-1object LEC vs. CA1: sCsO-sCdO of discriminating context-coding neurons | Wilcoxon sum rank test | W = 8726 | 4.74E-05 | LEC: 64 neurons; CA1:95 neurons | 6 vs. 3 mice |

**Supplementary Table 3. Statistics from all tests performed in the Supplementary Figures.**

| **Comparison** | **Statistical test** | **Test value** | **p-value** | **neurons (n)** | **Mice (N)** |
| --- | --- | --- | --- | --- | --- |
| 2 box LEC: Abs firing rate change of discriminating neurons | Kruskal-Wallis test | H = 74.794; df = 3 | 4.01E-13 | 184 | 13 mice |
| W1 vs. W2 | Wilcoxon signed-rank tests with Bonferroni correction | 8933 | 1.00E+00 |  |  |
| W1 vs. A1 | Wilcoxon signed-rank tests with Bonferroni correction | 3029 | 2.14E-13 |  |  |
| W1 vs. A2 | Wilcoxon signed-rank tests with Bonferroni correction | 4691 | 7.80E-07 |  |  |
| W2 vs. A1 | Wilcoxon signed-rank tests with Bonferroni correction | 2652 | 3.39E-15 |  |  |
| W2 vs. A2 | Wilcoxon signed-rank tests with Bonferroni correction | 4312 | 3.93E-08 |  |  |
| A1 vs. A2 | Wilcoxon signed-rank tests with Bonferroni correction | 9997 | 2.39E-01 |  |  |
| 2 room LEC: Abs firing rate change of discriminating neurons | Kruskal-Wallis test | H = 20.673; df = 3 | 0.0001231 | 75 neurons | 5 mice |
| W1 vs. W2 | Wilcoxon signed-rank tests with Bonferroni correction | 1198 | 1 |  |  |
| W1 vs. A1 | Wilcoxon signed-rank tests with Bonferroni correction | 622 | 1.36E-04 |  |  |
| W1 vs. A2 | Wilcoxon signed-rank tests with Bonferroni correction | 660 | 0.000325 |  |  |
| W2 vs. A1 | Wilcoxon signed-rank tests with Bonferroni correction | 768 | 0.003 |  |  |
| W2 vs. A2 | Wilcoxon signed-rank tests with Bonferroni correction | 756 | 0.002 |  |  |
| A1 vs. A2 | Wilcoxon signed-rank tests with Bonferroni correction | 1607 | 1 |  |  |
| 2 box CA1: Abs firing rate change of discriminating neurons | Kruskal-Wallis test | H = 25.696; df = 3 | 1.10E-05 | 167 neurons | 4 mice |
| W1 vs. W2 | Wilcoxon signed-rank tests with Bonferroni correction | 6886 | 1 |  |  |
| W1 vs. A1 | Wilcoxon signed-rank tests with Bonferroni correction | 4166 | 3.22E-05 |  |  |
| W1 vs. A2 | Wilcoxon signed-rank tests with Bonferroni correction | 4491 | 3.34E-04 |  |  |
| W2 vs. A1 | Wilcoxon signed-rank tests with Bonferroni correction | 4134 | 2.52E-05 |  |  |
| W2 vs. A2 | Wilcoxon signed-rank tests with Bonferroni correction | 4354 | 1.28E-04 |  |  |
| A1 vs. A2 | Wilcoxon signed-rank tests with Bonferroni correction | 7113 | 1.00E+00 |  |  |
| 2 room CA1: Abs firing rate change of discriminating neurons | Kruskal-Wallis test | H = 62.157; df = 3 | 2.03E-10 | 128 neurons | 4 mice |
| W1 vs. W2 | Wilcoxon signed-rank tests with Bonferroni correction | 5137 | 9.90E-02 |  |  |
| W1 vs. A1 | Wilcoxon signed-rank tests with Bonferroni correction | 2042 | 4.24E-06 |  |  |
| W1 vs. A2 | Wilcoxon signed-rank tests with Bonferroni correction | 2231 | 3.89E-05 |  |  |
| W2 vs. A1 | Wilcoxon signed-rank tests with Bonferroni correction | 1464 | 1.43E-09 |  |  |
| W2 vs. A2 | Wilcoxon signed-rank tests with Bonferroni correction | 1857 | 4.01E-07 |  |  |
| A1 vs. A2 | Wilcoxon signed-rank tests with Bonferroni correction | 4489 | 1.00E+00 |  |  |
| 2 box vs. 2 room LEC FR change | Wilcoxon sum rank test | W = 6731 | 7.58E-01 |  |  |
| 2 box vs. 2 room CA1 FR change | Wilcoxon sum rank test | W = 13133 | 0.0007614 |  |  |
| 2-box LEC 5718: map stability of spatial neurons | Kruskal-Wallis test | H = 35.422; df = 3 | 9.92E-05 | 31 neurons | 1 |
| W1 vs. W2 | Wilcoxon signed-rank tests with Bonferroni correction | 289 | 1.00E+00 |  |  |
| W1 vs. A1 | Wilcoxon signed-rank tests with Bonferroni correction | 450 | 1.04E-04 |  |  |
| W1 vs. A2 | Wilcoxon signed-rank tests with Bonferroni correction | 459 | 3.19E-05 |  |  |
| W2 vs. A1 | Wilcoxon signed-rank tests with Bonferroni correction | 434 | 6.42E-04 |  |  |
| W2 vs. A2 | Wilcoxon signed-rank tests with Bonferroni correction | 482 | 6.12E-07 |  |  |
| A1 vs. A2 | Wilcoxon signed-rank tests with Bonferroni correction | 304 | 1 |  |  |
| 2-room LEC 5718: map stability of spatial neurons | Kruskal-Wallis test | H = 77.056; df = 3 | < 2.2e-16 | 74 | 1 |
| W1 vs. W2 | Wilcoxon signed-rank tests with Bonferroni correction | 1302 | 1 |  |  |
| W1 vs. A1 | Wilcoxon signed-rank tests with Bonferroni correction | 2587 | 6.30E-10 |  |  |
| W1 vs. A2 | Wilcoxon signed-rank tests with Bonferroni correction | 2622 | 1.78E-10 |  |  |
| W2 vs. A1 | Wilcoxon signed-rank tests with Bonferroni correction | 2581 | 7.80E-10 |  |  |
| W2 vs. A2 | Wilcoxon signed-rank tests with Bonferroni correction | 2549 | 2.39E-09 |  |  |
| A1 vs. A2 | Wilcoxon signed-rank tests with Bonferroni correction | 1333 | 1 |  |  |
| 2-box LEC 5721: map stability of spatial neurons | Kruskal-Wallis test | H = 86.004; df = 3 | < 2.2e-16 | 63 | 1 |
| W1 vs. W2 | Wilcoxon signed-rank tests with Bonferroni correction | 1299 | 2.80E-01 |  |  |
| W1 vs. A1 | Wilcoxon signed-rank tests with Bonferroni correction | 1887 | 1.09E-08 |  |  |
| W1 vs. A2 | Wilcoxon signed-rank tests with Bonferroni correction | 1870 | 2.21E-08 |  |  |
| W2 vs. A1 | Wilcoxon signed-rank tests with Bonferroni correction | 1840 | 7.50E-08 |  |  |
| W2 vs. A2 | Wilcoxon signed-rank tests with Bonferroni correction | 1972 | 2.53E-10 |  |  |
| A1 vs. A2 | Wilcoxon signed-rank tests with Bonferroni correction | 1009 | 1 |  |  |
| 2-room LEC 5721: map stability of spatial neurons | Kruskal-Wallis test | H = 64.981; df = 3 | 5.06E-11 | 48 neurons | 1 |
| W1 vs. W2 | Wilcoxon signed-rank tests with Bonferroni correction | 643 | 1.00E+00 |  |  |
| W1 vs. A1 | Wilcoxon signed-rank tests with Bonferroni correction | 1141 | 1.84E-10 |  |  |
| W1 vs. A2 | Wilcoxon signed-rank tests with Bonferroni correction | 1158 | 1.08E-10 |  |  |
| W2 vs. A1 | Wilcoxon signed-rank tests with Bonferroni correction | 1144 | 1.18E-10 |  |  |
| W2 vs. A2 | Wilcoxon signed-rank tests with Bonferroni correction | 1137 | 3.23E-10 |  |  |
| A1 vs. A2 | Wilcoxon signed-rank tests with Bonferroni correction | 658 | 1 |  |  |
| LEC A1 map stability 2box vs. 2room vs. 2 ctrl | Kruskal-Wallis test | H = 9.6805; df = 2 | 0.007905 | 2box: 307; 2 room: 189; 2ctrl: 22 | 2box: 13; 2 room: 5; 2ctrl: 4 |
| 2box vs. 2room | Tukey multiple comparisons of means | diff = -0.03501326 | 2.93E-01 |  |  |
| 2box vs. 2ctrl | Tukey multiple comparisons of means | diff = 0.12974889 | 4.65E-02 |  |  |
| 2room vs. 2ctrl | Tukey multiple comparisons of means | diff = 0.16478215 | 5.55E-03 |  |  |
| CA1 A1 map stability 2box vs. 2room vs. 2 ctrl | Kruskal-Wallis test | H = 88.929; df = 2 | < 2.2e-16 | 2box: 212; 2 room: 181; 2ctrl: 52 | 2box: 4; 2 room: 4; 2ctrl: 4 |
| 2box vs. 2room | Tukey multiple comparisons of means | 0.009175852 | 0.9459619 |  |  |
| 2box vs. 2ctrl | Tukey multiple comparisons of means | 0.504414855 | < 2.2e-16 |  |  |
| 2room vs. 2ctrl | Tukey multiple comparisons of means | 0.495239003 | < 2.2e-16 |  |  |
| 2-room CA1: map stability of FS spatial neurons | Kruskal-Wallis test | H = 57.542; df = 3 | 0 | 25 | 4 |
| W1 vs. W2 | Wilcoxon signed-rank tests with Bonferroni correction | 114 | 1 |  |  |
| W1 vs. A1 | Wilcoxon signed-rank tests with Bonferroni correction | 325 | 3.58E-07 |  |  |
| W1 vs. A2 | Wilcoxon signed-rank tests with Bonferroni correction | 324 | 7.14E-07 |  |  |
| W2 vs. A1 | Wilcoxon signed-rank tests with Bonferroni correction | 320 | 3.58E-06 |  |  |
| W2 vs. A2 | Wilcoxon signed-rank tests with Bonferroni correction | 320 | 3.58E-06 |  |  |
| A1 vs. A2 | Wilcoxon signed-rank tests with Bonferroni correction | 155 | 1.00E+00 |  |  |
| 2-room LEC: map stability of FS spatial neurons | Kruskal-Wallis test | H = 9.6126; df = 3 | 2.22E-02 | 9 | 5 |
| W1 vs. W2 | Wilcoxon signed-rank tests with Bonferroni correction | 26 | 1.00E+00 |  |  |
| W1 vs. A1 | Wilcoxon signed-rank tests with Bonferroni correction | 45 | 0.023 |  |  |
| W1 vs. A2 | Wilcoxon signed-rank tests with Bonferroni correction | 45 | 0.023 |  |  |
| W2 vs. A1 | Wilcoxon signed-rank tests with Bonferroni correction | 40 | 2.35E-01 |  |  |
| W2 vs. A2 | Wilcoxon signed-rank tests with Bonferroni correction | 45 | 2.30E-02 |  |  |
| A1 vs. A2 | Wilcoxon signed-rank tests with Bonferroni correction | 34 | 1.00E+00 |  |  |
